# Population genomic evidence that human and animal infections in Africa come from the same populations of *Dracunculus medinensis*

**DOI:** 10.1101/808923

**Authors:** Caroline Durrant, Elizabeth A. Thiele, Nancy Holroyd, Stephen R Doyle, Guillaume Sallé, Alan Tracey, Geetha Sankaranaranayan, Magda E. Lotkowska, Hayley M. Bennett, Thomas Huckvale, Zahra Abdellah, Ouakou Tchindebet, Mesfin Wossen, Makoy Samuel Yibi Logora, Cheick Oumar Coulibaly, Adam Weiss, Albrecht I Schulte-Hostedde, Jeremy Foster, Christopher A. Cleveland, Michael J. Yabsley, Ernesto Ruiz-Tiben, Matthew Berriman, Mark L. Eberhard, James A. Cotton

**Affiliations:** Wellcome Sanger Institute, Hinxton, Cambridgeshire, United Kingdom; Department of Biology, Vassar College, Poughkeepsie, New York, USA; INRA - U. Tours, UMR 1282 ISP Infectiologie et Santé Publique, Nouzilly, France; The Carter Center, Atlanta, Georgia, USA; Department of Biology, Laurentian University, Sudbury, Canada; New England Biolabs, Ipswich, Massachusetts, USA; University of Georgia, Athens, Georgia, USA; Centers for Disease Control and Prevention, Atlanta, Georgia, USA; Berkeley Lights Inc., Emeryville, California, USA

## Abstract

**Background:** Guinea worm – *Dracunculus medinensis* – was historically one of the major parasites of humans and has been known since antiquity. Now, Guinea worm is on the brink of eradication, as efforts to interrupt transmission have reduced the annual burden of disease from millions of infections per year in the 1980s to only 30 human cases reported globally last year. Despite the enormous success of eradication efforts to date, one complication has arisen. Over the last few years, hundreds of dogs have been found infected with this previously apparently anthroponotic parasite, almost all in Chad. Moreover, the relative numbers of infections in humans and dogs suggests that dogs may be key in maintaining transmission in that country.

**Results:** In an effort to shed light on this peculiar epidemiology of Guinea worm in Chad, we have sequenced and compared the genomes of worms from dog, human and other animal infections. Confirming previous work with other molecular markers, we show that all of these worms are *D. medinensis*, and that the same population of worms are causing both infections, can confirm the suspected transmission between host species and detect signs of a population bottleneck due to the eradication efforts. The diversity of worms in Chad appears to exclude the possibility that there were no, or very few, worms present in the country during a 10-year absence of reported cases.

**Conclusions:** This work reinforces the importance of adequate surveillance of both human and dog populations in the Guinea worm eradication campaign and suggests that control programs should stay aware of the possible emergence of unusual epidemiology as they approach elimination.

## Background

Guinea worm – *Dracunculus medinensis* (Linnaeus, 1758) Gallandant, 1773 – has been an important human parasite for most of history. It is also one of the best known human pathogens and has been known since antiquity (Muller, 1971). This infamy is presumably due to its distinctive life cycle, where the large adult female worm (up to 1m long) causes excruciating pain as it emerges from a skin lesion. As recently as 1986, there were probably over 3 million cases of Guinea worm disease (GWD) from 22 countries in Africa and Asia (Watts, 1987) and historically probably very many more (Stoll, 1947). Called the quintessential “forgotten disease of forgotten people,” GWD was responsible for an enormous disease burden as patients are incapacitated for several weeks during worm emergence (Weiss et al., 2018; and many other studies cited in Ruiz-Tiben & Hopkins, 2006), and subsequent complications and serious secondary infections of the resulting ulcer are common and occasionally fatal (Muller, 1971).

Following the eradication of smallpox in 1980, public health scientists at the US Centers for Disease Control and Prevention (CDC) recognised that Guinea worm disease was a potential target for eradication (e.g. Muller, 1979; Bourne, 1982). Since 1986, Guinea worm has been the target of a large-scale control program aiming for complete, global eradication of the disease and extinction of the parasite responsible (Cairncross, Muller, & Zagaria, 2002). The introduction of interventions to encourage residents to report cases of GWD, prevent infected persons from contaminating source of drinking water, provide new sources of safe water and promote greater use of existing sources, promote the use of cloth and pipe (“straw”) filters, and the application of vector control measures has subsequently reduced the incidence of GWD (Hopkins & Ruiz-Tiben, 1991; Ruiz-Tiben & Hopkins, 2006). As the program has progressed, these measures have been complemented by work to ensure sources of infection are traced and treated for cases, containment of cases to prevent contamination of water, and active searching for new cases (Hopkins & Ruiz-Tiben, 1991). The Guinea worm eradication program has been a great success – by 2000 there were only 74,258 cases of GWD in 15 countries in sub-Sharan Africa (Centers for Disease Control and Prevention, 2001), and this had fallen to just 15 cases in each of Chad and Ethiopia as of 2017 (Hopkins et al., 2018). Either Guinea worm disease or polio (Grassly & Orenstein, 2018) will soon become the second human disease to be eradicated, and Guinea worm is on track to be the first to be wiped out without a vaccine, and probably the first animal species to be deliberately made extinct.

The eradication campaign was predicated on *D. medinensis* being an anthroponotic parasite, with transmission between people via drinking water. Sporadic reports of animal infections were assumed to either be due to misidentification of the worm involved or to represent spill-over infection of little or no epidemiological importance. However, experimental infections of non-human animals – and particularly of dogs – have been performed successfully on a number of occasions (Muller, 1971). In natural conditions, worms have particularly frequently been reported as emerging from dogs, but generally at a low prevalence and sporadically. When human infections with Guinea worm have been eliminated from a region, dog infections from that region have subsequently disappeared (Eberhard, Ruiz-Tiben, & Hopkins, 2016; Eberhard et al., 2014). There are some apparent exceptions: for example, in Bukhara, Uzbekistan, where a hotspot of very high Guinea worm prevalence (up to 20%) was eliminated in the 1930s, Guinea worm infections in dogs continued to be reported, but no human cases were found after 1931 (World Health Organisation, 1998; Litvinov & Lysenko, 1985).

From 2012, however, a distinct, and apparently unique situation became evident in Chad, where large numbers of infections in domestic dogs have appeared, against the background of a small number of human cases (Eberhard et al., 2014). Dog infections became evident beginning in April 2012, when the Chad Guinea worm eradication program (with assistance from The Carter Center) launched active village-based surveillance in nearly 700 villages, following the detection in 2010 of human cases for the first time in 10 years in Chad. With increasing surveillance of dog populations, the number of dog infections reported has subsequently steadily increased, and in 2016, there were over 1,000 infected dogs reported from Chad. Small numbers of dog infections have also been identified in the other recently endemic countries (in 2016, 14 from Ethiopia, 11 from Mali and none from South Sudan, although this country did report a single dog infection in 2015). Greater scrutiny of animals for potential Guinea worm infection has also revealed occasional infections in wildlife, such as cats and baboons (see Hopkins, Ruiz-Tiben, Eberhard, Roy, & Weiss, 2017 for a full description of the situation in 2016-2017).

In this context, there was some uncertainty as to whether the worms emerging from dog and human infections in Chad represented the same species. Most of the key defining morphological features for this group of nematodes are found on adult males, which are not recovered from natural infections (Muller, 1971; Cleveland et al., 2018). The other described species of the genus *Dracunculus* are all from the New World and include *D. insignis* and *D. lutrae* from North American carnivorous mammals and *D. fuelleborni* from a Brazilian opossum (Jones & Mulder, 2007; Muller, 1971; Travassos, 1934). There are also numerous reports of *Dracunculus* spp. in reptiles – particularly snakes (see Cleveland et al., 2018). Molecular phylogenetic work supports the mammalian parasites as a distinct clade to those found in other vertebrates (Wijova, Moravec, Horak, Modry, & Lukes, 2005; Bimi, Freeman, Eberhard, Ruiz-Tiben, & Pieniazek, 2005; Elsasser, Floyd, Hebert, & Schulte-Hostedde, 2009; Nadler et al., 2007). The diversity and phylogeny of the genus has recently been reviewed (Cleveland et al., 2018). There is a relative scarcity of parasitological work on wild mammals, and particularly of work looking beyond gastrointestinal species. There are also a number of reports of of cryptic species of parasitic worms in wildlife (reviewed in Cole & Viney, 2018). It is thus possible that other, undescribed mammal-infective species exist, and these could explain the few reports of human or mammal Guinea worm infections from countries otherwise considered non-endemic (see Muller, 1971 for references to case reports). A number of comprehensive reviews of *D. medinensis* biology, epidemiology and control are available (Muller, 1971; e.g. Cairncross et al., 2002) and the reader is referred to the extensive literature on the Guinea worm eradication program (Hopkins & Ruiz-Tiben, 1991; Ruiz-Tiben & Hopkins, 2006; Biswas, Sankara, Agua-Agum, & Maiga, 2013; e.g. Al-Awadi et al., 2014; Molyneux & Sankara, 2017), including regularly updated surveillance data (most recently in Hopkins et al., 2018).

Previous molecular work established that *D. medinensis* and *D. insignis* could be differentiated at the *18S rRNA* locus, and that a single dog worm from Ghana was identical at that locus to *D. medinensis* collected from nearby human cases (Bimi et al., 2005). We subsequently reported data from the *18S rRNA* locus and a mitochondrial marker for 14 worms that emerged from humans and 17 from dogs in Chad, together with whole-genome data from 6 worms (Eberhard et al., 2014). A draft reference genome assembly based on sequence data from a single worm from Ghana (International Helminth Genomes Consortium, 2017) recently gave the first picture of the genome content of this species and confirmed the phylogenetic position of *D. medinensis* within a large spiruromorph clade of parasites related to filarial nematodes. Here, we present an improved genome assembly for *D. medinensis* and whole genome sequence data for a much larger set of adult *D. medinensis* and from two closely related species. Together, these data give a detailed picture of the relationships between *D. medinensis* from different hosts and countries, and confirms existing microsatellite genotyping and mitochondrial sequence data from the same populations (Thiele et al., 2018) which showed that human cases of dracunculiasis and animal infections all originate from the same populations of *D. medinensis*.

## Results

### Whole-genome sequence data from *Dracunculus* specimens from a range of host species and geographic regions

In this study, we attempted to generate whole-genome shotgun sequence data for 90 *D. medinensis* specimens; for four samples we could not make sequencing libraries. We also sequenced two samples of *D. insignis* and one sample of *D. lutrae*. To aid interpretation of these data, we used the original Illumina data used to improve the v2.0.4 reference genome assembly for *D. medinensis* – based on a worm collected in Ghana in 2001 (International Helminth Genomes Consortium, 2017) – with a combination of both manual and automated approaches to produce an improved (v3.0) assembly for *D. medinensis.* This substantially improved contiguity and reduced misassemblies, for example the average scaffold length is twice that of the previously published assembly version (Supplementary table 1).

Despite extensive sequencing effort, mapping our data against this reference showed that we achieved a median depth of 10x coverage for only about one-third (33) of *D. medinensis* samples. Unless otherwise specified, subsequent analyses were restricted to this set of 33 *D. medinensis* samples. These samples were collected from a number of African countries (Figure 1), with 22 from Chad, 5 from Ethiopia, 2 each from Ghana and South Sudan and 1 each from Mali and Côte d’Ivoire. It included 15 samples from humans, 15 from dogs and 3 from other animals (2 Ethiopian baboons, *<u>Papio anubis</u>*, and one from a Chad cat, *Felis catus*). Full details of the samples and coverage achieved are shown in Table 1. The low coverage we achieved was due to extensive contamination with bacterial and, in some cases, host DNA, so that the percentage of reads mapping to the reference varied from 0.07% up to 94.8% (see Figure 1; Supplementary Figure 1); even within the genome-wide coverage set of 33 samples as few as 17.9% of reads mapped in one case. For 9 *D. medinensis* samples, sequence libraries were generated from both adult female material and L1 larvae present in the same sample tubes (representing the offspring of that female).

**Figure 1.**
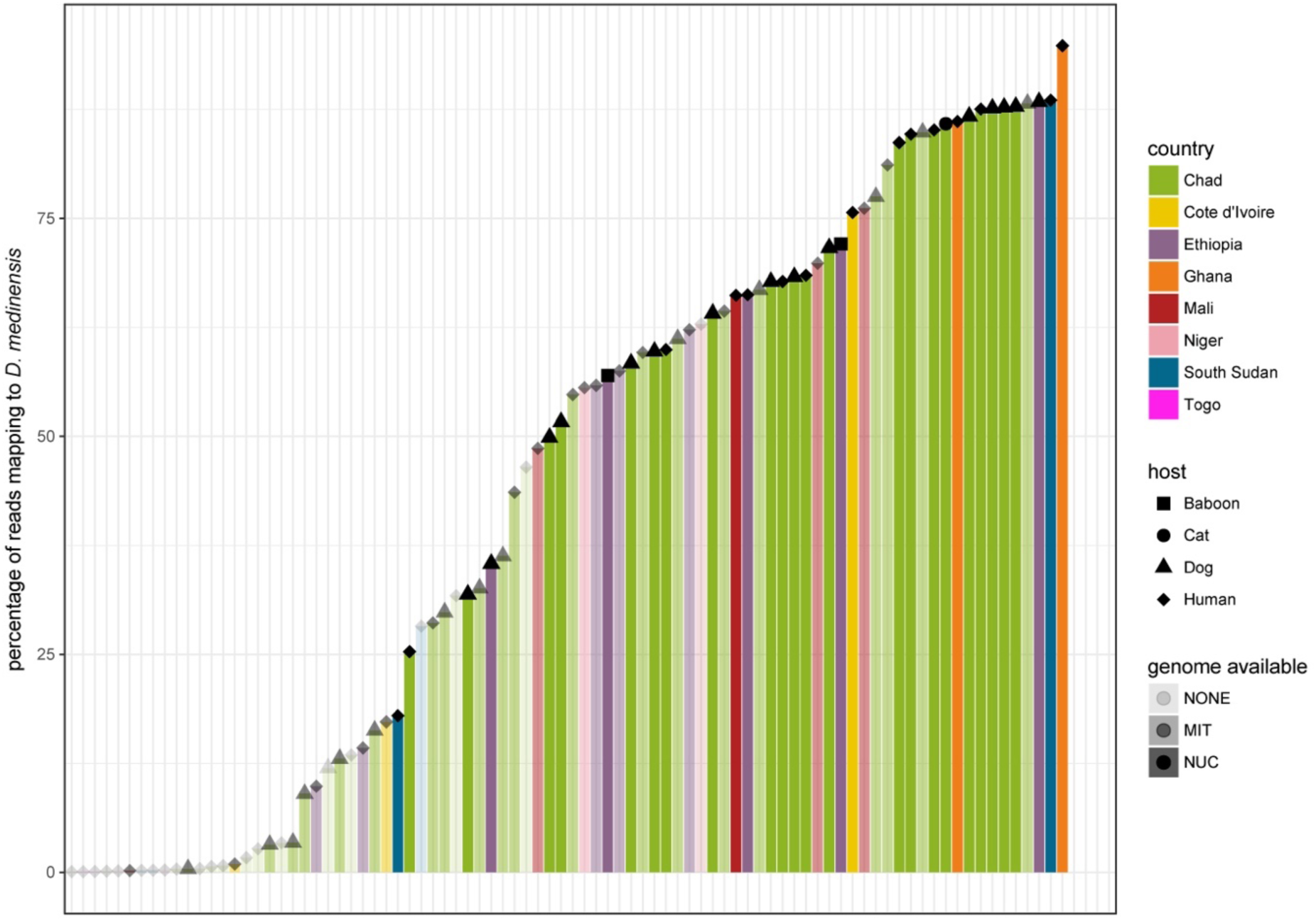
Proportion of sequencing reads mapping to the reference genome assembly for *Dracunculus medinensis* samples. Each bar indicates the proportion of sequencing reads from each sample that mapped against the reference genome assembly. The density of each bar indicates whether whole-genome data is included in our analysis, only mitochondrial genome data or whether insufficient data was available for that sample.

**Table 1.** Assembly statistics for *Dracunculus medinensis* assembly versions

|  | *D. medinensis* v2.0.4 | *D. medinensis* v3.0 |
| --- | --- | --- |
| **Total length (bp)** | 103,750,892 | 103,601,578 |
| **Number of scaffolds** | 1350 | 672 |
| **Average scaffold length (bp)** | 76,853 | 154,169 |
| **N50 scaffold length (bp)** | 665,026 | 3,396,158 |
| **Number of scaffolds > N50** | 33 | 10 |
| **N90 scaffold length (bp)** | 74,011 | 374,449 |
| **Number of scaffolds > N90** | 240 | 42 |
| **Total gap length (bp)** | 167,953 | 38,232 |

**Table 2.** Population genetic summary statistics for *Dracunculus medinensis* populations. Values are means and 95% bootstrap confidence intervals for the means of 1kb windows containing between 5 and 100 informative (variable) sites.

| population | $\Pi$ | Watterson's $\theta$ | Tajima's D |
| --- | --- | --- | --- |
| Chad | 0.0252<br>(0.0244- 0.0259) | 0.0130<br>(0.0126,0.0135) | 0.0637<br>(0.0617,0.0658) |
| East Africa | 0.0217<br>(0.0209- 0.0225) | 0.0154<br>(0.0148,0.0159) | 0.0410<br>(0.0392,0.0429) |
| West Africa | 0.00126<br>(0.000973-0.00162) | 0.00118<br>(0.000892,0.001508) | -0.00154<br>(-0.00308,-0.000139) |

Coverage also varied across the genome, most strikingly for two of the longest 5 scaffolds, which were often lower in coverage than other large scaffolds, varying from around three-quarters of the expected coverage to approximately similar coverage. We hypothesised that these scaffolds could represent all or part of a sex chromosome (X) in *D. medinensis*. The L1 larval samples showed coverage on these scaffolds around 75% that for other large scaffolds, as expected for an XY or XO sex determination system if the pool of larvae consisted of an approximately equal ratio of male and female larvae. More surprisingly, most of the DNA samples extracted from female worms showed similarly low relative coverage of these two scaffolds. We suggest that this is because much of the material extracted from these specimens is actually from L1 larvae remaining in the body of the female worm section. This seems plausible, as much of the female body comprises uterus containing several million larvae (Cairncross et al., 2002), and if the female body was largely degraded that explains the difficulty we had in extracting DNA from many samples.

To confirm this, we generated sequence data for juvenile worms harvested from a domestic ferret experimentally infected with *D. medinensis* (Eberhard, Yabsley, et al., 2016). These worms were 83 days old and pre-patent (and thus comprised only or largely somatic tissue), but could be morphologically identified as male and female. Analysis of data from these worms (Figure 2a) confirmed that the scaffolds with low coverage showed this pattern specifically in male worms, while the female worm showed essentially even coverage across the largest scaffolds, including the putative X and likely autosomal scaffolds. Further evidence comes from a comparison with *Onchocerca volvulus*, in which the sex chromosomes are known, as there is clear synteny between the *D. medinensis* scaffolds with variable coverage and one end of the *O. volvulus* X chromosome (Supplementary Figure 2). This part of the *O. volvulus* X chromosome represents the ancestral X chromosome of filarial nematode (Cotton et al., 2016): these data suggest that this was already present in *Dracunculus*, as well as filarial nematodes as previously suggested (Post, 2005).

**Figure 2.**
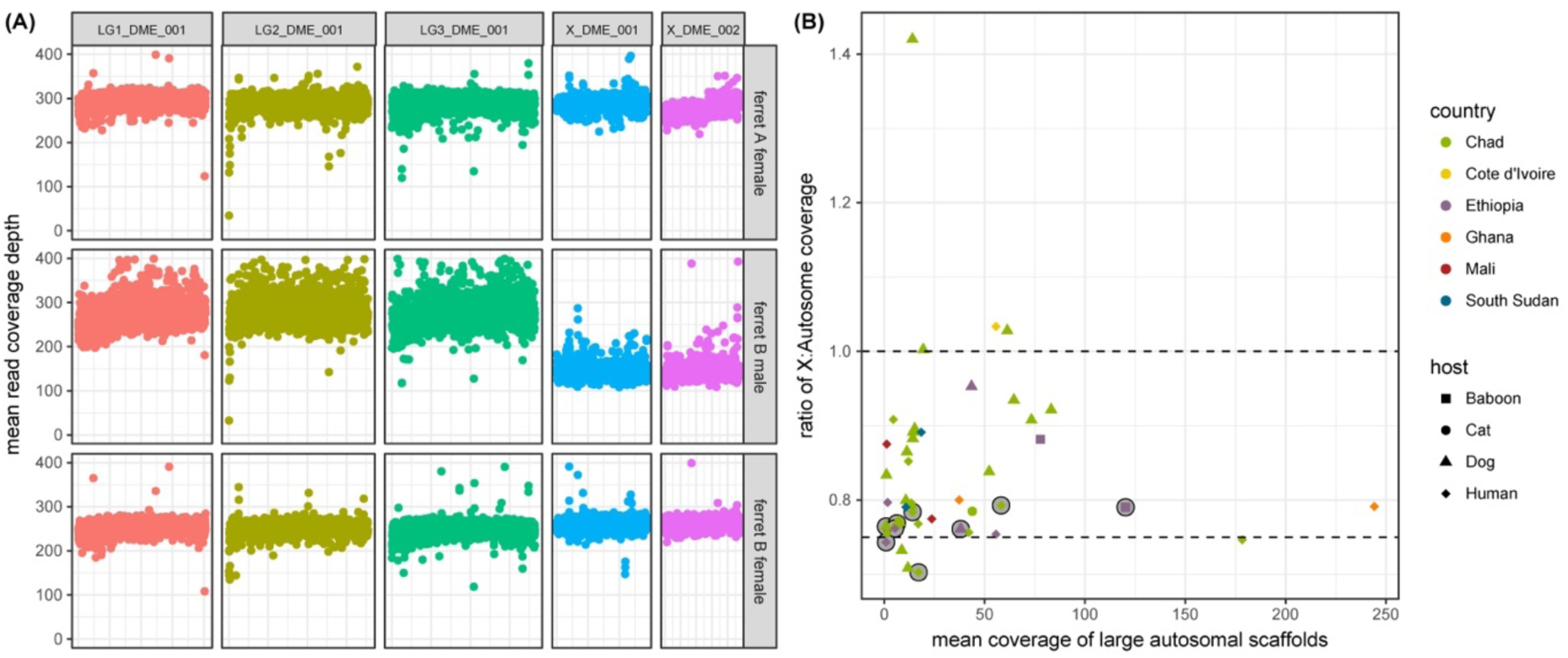
**(a)** Coverage variation across the *Dracunculus medinensis* genome in worms with known sex. Each point is the mean single read coverage across non-overlapping 5kb windows along the length of the five longest scaffolds for three juvenile worms recovered from an experimentally infected ferret. The 3 longest scaffolds show synteny to different *Onchocerca volvulus* chromosomes, the next 2 scaffolds are syntenic to the *O. volvulus* X chromosome (see Supplementary Figure S1). **(b**) Ratio of coverage across large autosomal and X-linked scaffolds for worm recovered from infected humans and animals in Africa. The y-axis shows the ratio of mean coverage on the 3 longest autosomal scaffolds to that of the mean coverage on the 2 longest X-linked scaffolds (these are the longest 5 scaffolds in the assembly, as shown in panel (a).

We thus used the ratio of mean coverage between the 3 largest autosomal scaffolds and 2 longest X chromosome scaffolds as a measure of the proportion of genomic DNA in our sample derived from larval vs female tissue, under the assumption that larvae are an equal mixture of the two sexes (Figure 2b). These data confirm that many samples contain substantial amounts of larval-derived DNA. One sample had a particularly high value for this statistic – for this sample, the mean coverage on scaffold X_DME_002 was inflated by the presence of a small region of extremely high read depth. X chromosome scaffolds were also excluded from subsequent population genetic analyses likely to be sensitive to the different dosage of these chromosomes (see Methods).

### African *D. medinensis* is highly divergent from other mammalian *Dracunculus* species

While most sequencing reads from high-quality *D. medinensis* samples mapped against our reference assembly (median across samples of 68.48%), the reads from the other two *Dracunculus* species mapped less comprehensively against the *D. medinensis* reference (Supplementary Table 1), which given the mapping parameters used suggests that many regions of the genome are more than 5% divergent between species. This was confirmed by variant calls in those regions of good read mapping: even given the poorer mapping quality, around 2.9 million sites varied between the three species, suggesting genome-wide divergence of at least 3% of the 103.8 Mb genome, as the mapping difficulty meant this is likely a significant underestimate. While interpretation of absolute divergence levels is difficult, our lower-bound estimate of divergence between these species is much greater than between different species of *Onchocerca* (*O. ochengi* and *O. volvulus*, respectively), which are less than 1% divergent (Cotton et al., 2016) and is consistent with having hundreds of thousands of years of independent evolutionary history. A principal components analysis (PCA) of SNP variants between these samples confirmed that samples from each species cluster closely together, and that the different species are well separated (Figure 3a). The first two principal component axes shown here explain 79.9% and 16.5% of the variation, respectively. More than 3-fold more sites were called as varying between *Dracunculus* spp. than observed across all 33 of our genome-wide *D. medinensis* samples, where about 981,198 sites vary.

**Figure 3.**
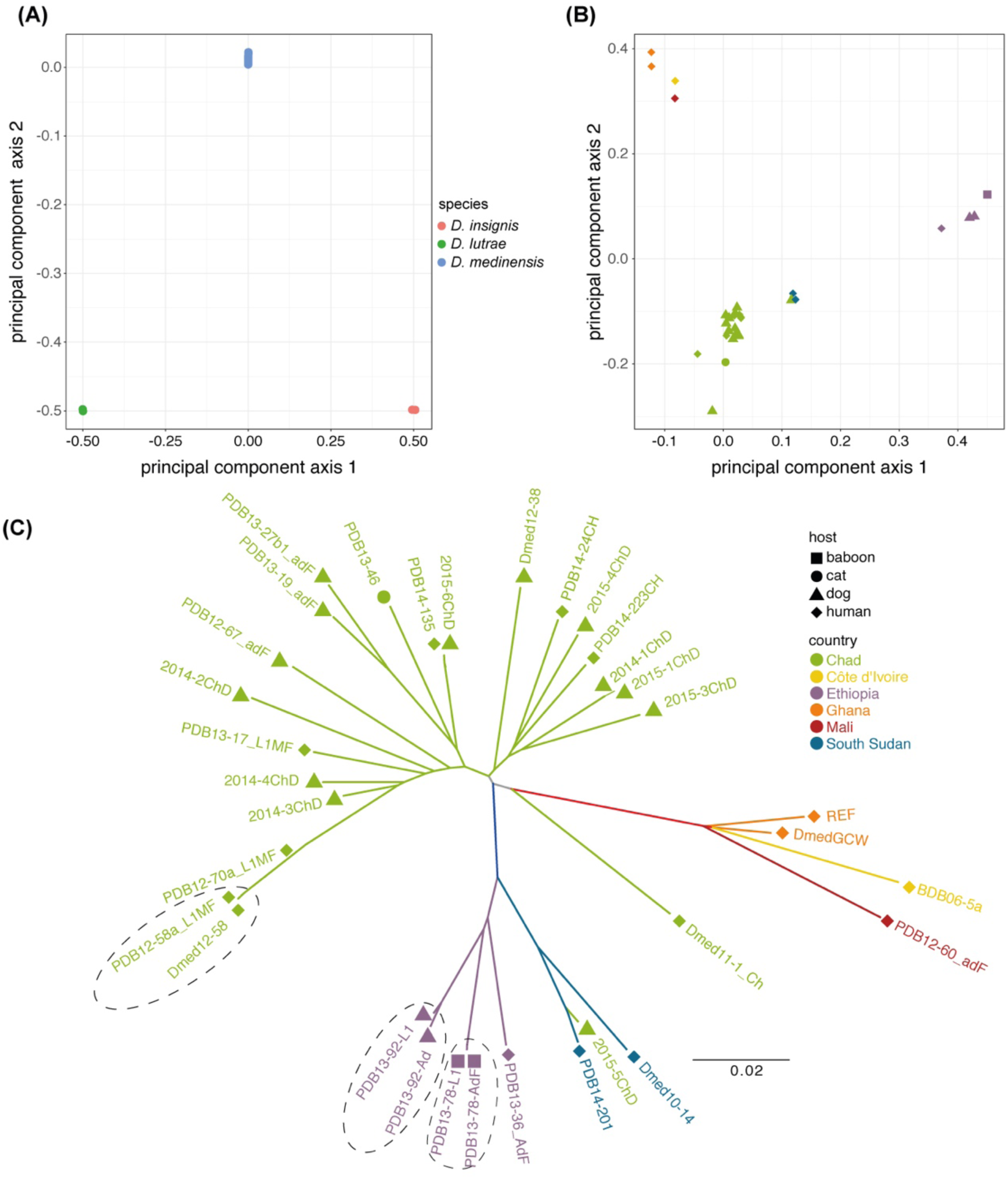
Principal components analysis of whole-genome data for **(a)** 33 *Dracunculus medinensis* samples, 2 *D. insignis* samples and 1 *D. lutrae* sample and **(b)** principal components analysis and **(c)** phylogenetic tree for just the 33 *D. medinensis* samples. The legend in the top right-hand corner of (c) applies to both panels (b) and (c). Dotted lines on panel (c) indicate three pairs of samples where both adult female tissue and L1 larvae from the same worm are included.

### Geography rather than host species explains the pattern of variation within African *D. medinensis*

Clear geographic structure was observed in the pattern of genome-wide variation within *D. medinensis*. PCA (Figure 3b) of the variants show distinct clusters of parasites from Ghana, Mali and Côte d’Ivoire (referred to as the ‘West African’ cluster) the Ethiopia, South Sudan and one Chad sample (an ‘East African’ cluster), and a group of parasites from Chad. The first two principal components explain only 22% of the variation in these data (14% and 8% respectively). Additional principal components axes, up to the 8^th^ axis, together explain 54% of the variation but none of these axes partition the genetic variation between host species (Supplementary Figure 3).

Phylogenetic analysis (Figure 3c) supported this pattern, with clear clades of West African and East African worms. The Chad sample visible as being part of the East African cluster in the PCA (2015-5ChD, a worm from a dog infection emerging in 2015) was part of the East African clade in the phylogenetic tree. A second Chad worm, from a human case in 2011, also appeared to be divergent from any other worm on our phylogeny. There was no apparent clustering by host species in Chad or Ethiopia, the two countries for which worms from multiple dog and human infections were included, and no clear clustering by year of worm emergence. In all three cases where both L1 larvae and adult sections from a single emerging worm yielded high-quality data, these two samples clustered very closely together.

Other approaches to investigate population structure support these conclusions. Bayesian clustering using MavericK strongly supported a model of only 2 populations (*K*=2) for these data, with posterior probability of 1.0 for this value of *K.* The two populations divided worms collected in Chad from those collected elsewhere, with the exception of the Chad worm 2015-5ChD, which clustered with those from other countries, as in the PCA and phylogeny. Analysis with Structure suggested that *K* values of between 2 and 4 fitted the data well. In all cases these analyses clustered worms largely by geographical origin, and not by host. In the highest K values, most Chad worms had mixed ancestry between two Chad populations, and in no analysis did worms from different host species cluster together more than expected (Supplementary Figure 4). As expected, differences in allele frequencies between worms from dog infections and human cases within Chad are low (mean F_st_ 0.01806, 99% confidence interval 0.0172-0.0189; median F_st_ 0.0114, CI 0.0109-0.0118) and consistently low across the genome (Supplementary Figure 5), confirming that there is no genetic difference between worms infecting dogs and humans.

**Figure 4.**
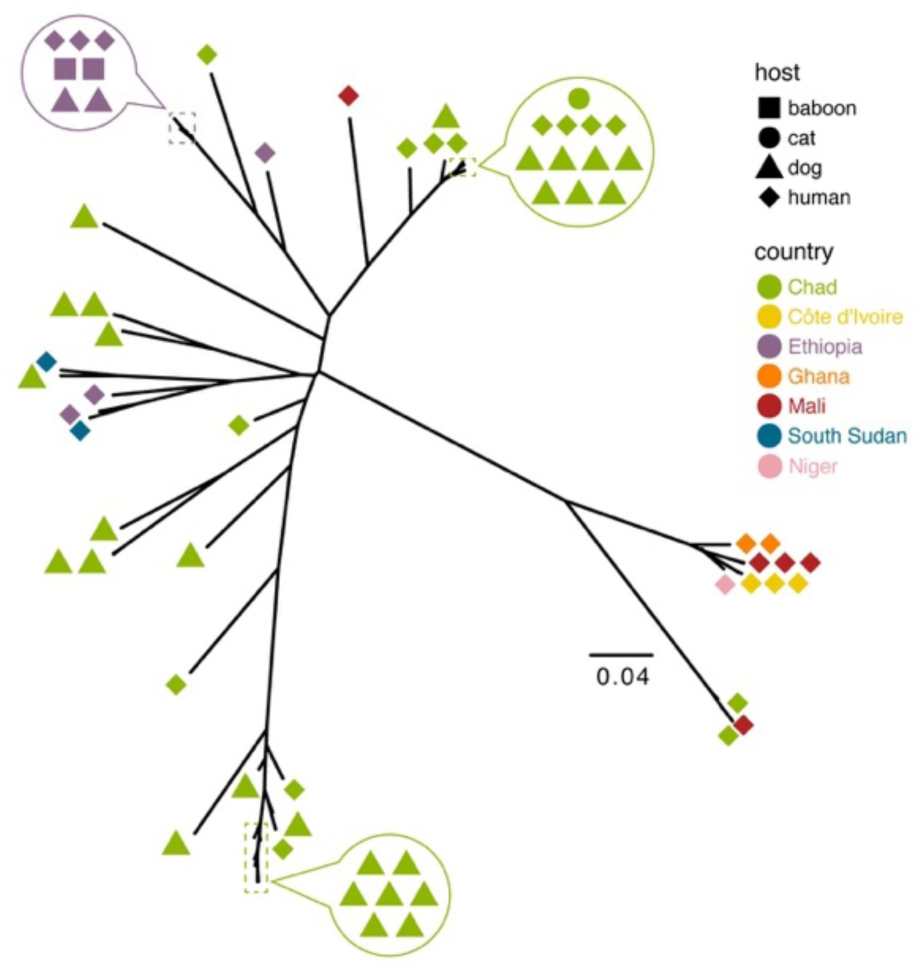
Phylogenetic tree based on inferred mitochondrial genome sequences for 65 *Dracunculus medinensis* samples for which sufficient coverage of the mitochondrial genome was available. For clarity, arrowed circles show host and geographic origin for samples with very similar mitochondrial haplotypes.

**Figure 5.**
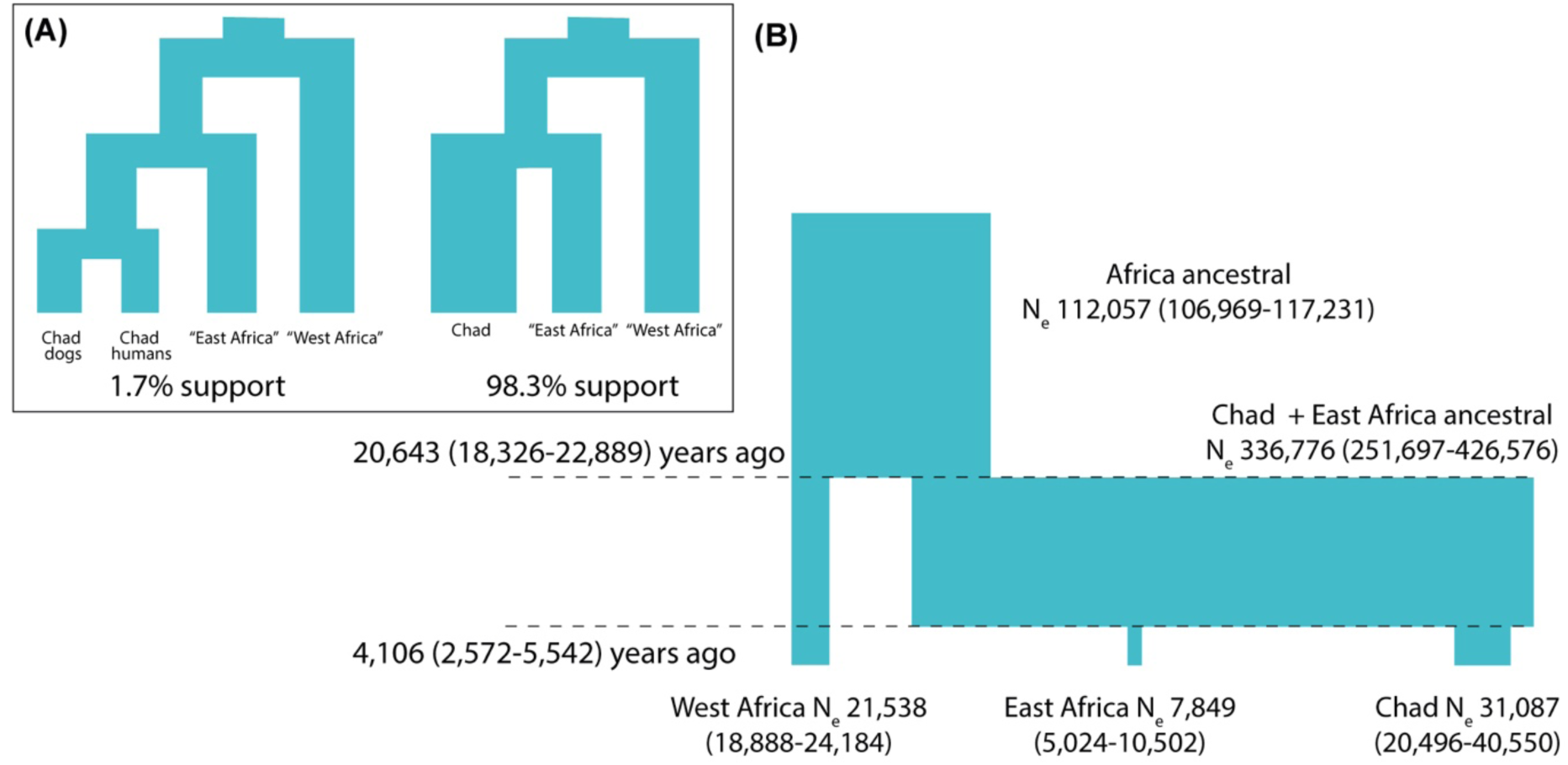
Coalescent models of *Dracunculus medinensis* population structure. (a) Out of all possible scenarios for up to 4 distinct isolated populations of *D. medinensis*, we find posterior support for only 2, with strong support only for a model in which all worms from Chad are part of one population, more closely related to worms from Ethiopia and South Sudan than to those from elsewhere in our sample set. (b) Estimates of divergence times and genetic (effective) population sizes under the supported model shown in (a). Values shown are posterior means and 95% highest posterior density estimates for each parameter in this model, under one set of prior assumptions.

### Mitochondrial genome data confirms the geographic structure of the *D. medinensis* population

To allow us to study a wider range of samples, we called variants against the mitochondrial genome of *D. medinensis* for a total of 65 samples that had median coverage of at least 10x across this sequence. The additional samples included 14 dog and 18 human samples and included a single sample from Niger, slightly expanding the geographical range of samples included. Our variant calling approach identified 182 variable sites that could be reliably genotyped across those samples. The results of this analysis (Figure 4) are congruent with those from nuclear genome variation, with a strong signal of clustering by geographical origin. The worm collected in Niger joined a tight cluster that included all West African samples (Ghana, Mali and Côte d’Ivoire) with the exception of two worms from Mali collected in 2014: one was closely related to two worms from Chad cases in 2014 and 2015, and the second appears as an outgroup to a large clade of Chad worms. The two other exceptions to the clear geographical structure were a worm from a dog in Chad in 2015 which was most similar to one from a South Sudan case from 2014 within a small clade of Ethiopia and South Sudan worms, and one from a human case in Chad in 2014 that groups as part of a more diverse group of Ethiopia worms. As with the nuclear data, worms from human cases and infections in dogs and other animals often group together, with extremely similar mitochondrial haplotypes; there is no clear signature of clustering by host species.

### *D. medinensis* from Chad are genetically diverse but are in decline

Phylogenetic analysis of both nuclear and mitochondrial data, and the nuclear genome PCA appear to show that worms in Chad are considerably more diverse than those from the other regions included in our analysis. To ensure an adequate sample size for comparison, we combined samples from countries with small numbers of samples into three regional groups, combining Ethiopia and South Sudan samples into an East African group, and samples from Mali, Ghana and Côte d’Ivoire into a West African group, while Chad was considered alone. Population genetic summary statistics (Supplementary Table 3) for these groups confirmed the pattern suggested by phylogenies and PCA: we see highest nucleotide diversity (Π) in Chad, while the East African group is slightly, but significantly less diverse and the West Africa group has an order-of-magnitude lower nucleotide diversity. A second estimator of genetic diversity (Watterson’s Θ) shows lower values for the East African and Chad populations, but is higher in East Africa than Chad. For neutral variants in a population at equilibrium Π and Θ are expected to be equal, but Watterson’s estimator is heavily influenced by rare alleles. The difference between Π and Θ that we observe indicates an excess of common variants in the East Africa and Chad populations over neutral expectations (see e.g. Charlesworth & Charlesworth, 2010 pp28-30 and pp288-289 for a full description). This is captured by high Tajima’s D values, which are simply a normalised difference between Π and Θ. While a variety of population genetic processes can influence these statistics, high Tajima’s D across the genome in these two regions is most likely indicative of a demographic process, such as a recent sharp decline in the worm populations (Tajima, 1989).

### Coalescent models suggest a large population has been continuously present in Chad

To confirm the population structure of *D. medinensis* in Africa, we constructed coalescent models based on 1kb loci spaced every 100kb – much longer than the distance over which linkage disequilibrium decays to approximately background levels – across the large scaffolds of the *D. medinensis* reference genome assembly. Due to our small number of samples, we combined samples into three regional groups. Ethiopia and South Sudan samples were combined into an East African group, samples from Mali, Ghana and Côte d’Ivoire into a West African group; and samples from Chad comprised their own group. Our more extensive sample of worms from Chad meant we could investigate whether Chad worms were best explained as two host-specific populations of worms from dog infections and human cases, or as a single group. Only two scenarios for the population structure received support in the posterior sample from the Markov chain Monte Carlo (MCMC) procedure (see Figure 5a). In both scenarios, worms from Chad were more closely related to those from the East African group than to the West African group. By far the strongest support (average 97.9% of posterior samples, over 3 replicate sets of 100 random loci) supported a single Chad population of worms that emerged from both human and dog hosts.

Using this highly supported population history, we used a second coalescence approach to estimate parameters describing the demographic history of the three regional present-day populations and the ancestral populations that gave rise to them (Figure 5b). Assuming a similar per-generation mutation rate to *C. elegans* and a generation time of 1 year, these analyses suggest that the Chad and East African populations have been separated for at least several thousand years, and that divergence from the West African population was about 5-fold older. The long-term effective population sizes of the Chad and East Africa populations reflect the higher nucleotide and phylogenetic diversity, with Chad being around 4-fold higher with an estimated 20 to 40 thousand breeding individuals.

### Relatedness between *D. medinensis* isolates

Our population genetic evidence supports the idea that a single, diverse population of Guinea worms exists in Chad and is infecting both humans and animal species. More direct evidence of transmission between host species would be genetic relatedness between worms that emerged in different species. We employed a method to estimate pairwise relatedness between isolates based on SNP variants that is intended to be robust to population structure. Kinship is the probability that a random allele sampled from each of two individuals at a particular locus are identical by descent. The expected value in an outbred diploid population is 0.5 for monozygotic twins and 0.25 for full sibs or parent-offspring pairs.

The median kinship across all pairs of samples we find is low, but non-zero (0.0078; approximately that expected for third cousins), but worms from the same countries are much more highly related (e.g. average kinship of 0.0846 for pairs within Chad). There is clear geographic structure to kinship in these data, as most worm samples from the same countries are related to at least one other sample from that country with kinship of close to 0.25 or higher (Figure 6), while only a single pair of closely related worms are from different countries. Notably, six pairs of worms have relatedness of higher than 0.45, close to the maximum possible value of 0.5 (Figure 6). These six pairs include all 3 sets of matched adult and larval samples in the whole-genome coverage set, providing support for the hypothesis that other pairs with a similarly high relatedness could represent either parent-offspring or full sibling pairs. The inflated kinship values is explained by a high level of inbreeding within each country. The other three pairs of high-relatedness samples are all from different worms and from consecutive years, so we interpret these as being parent-offspring pairs and these links thus represent putative direct transmission events between guinea worm infections.

**Figure 6.**
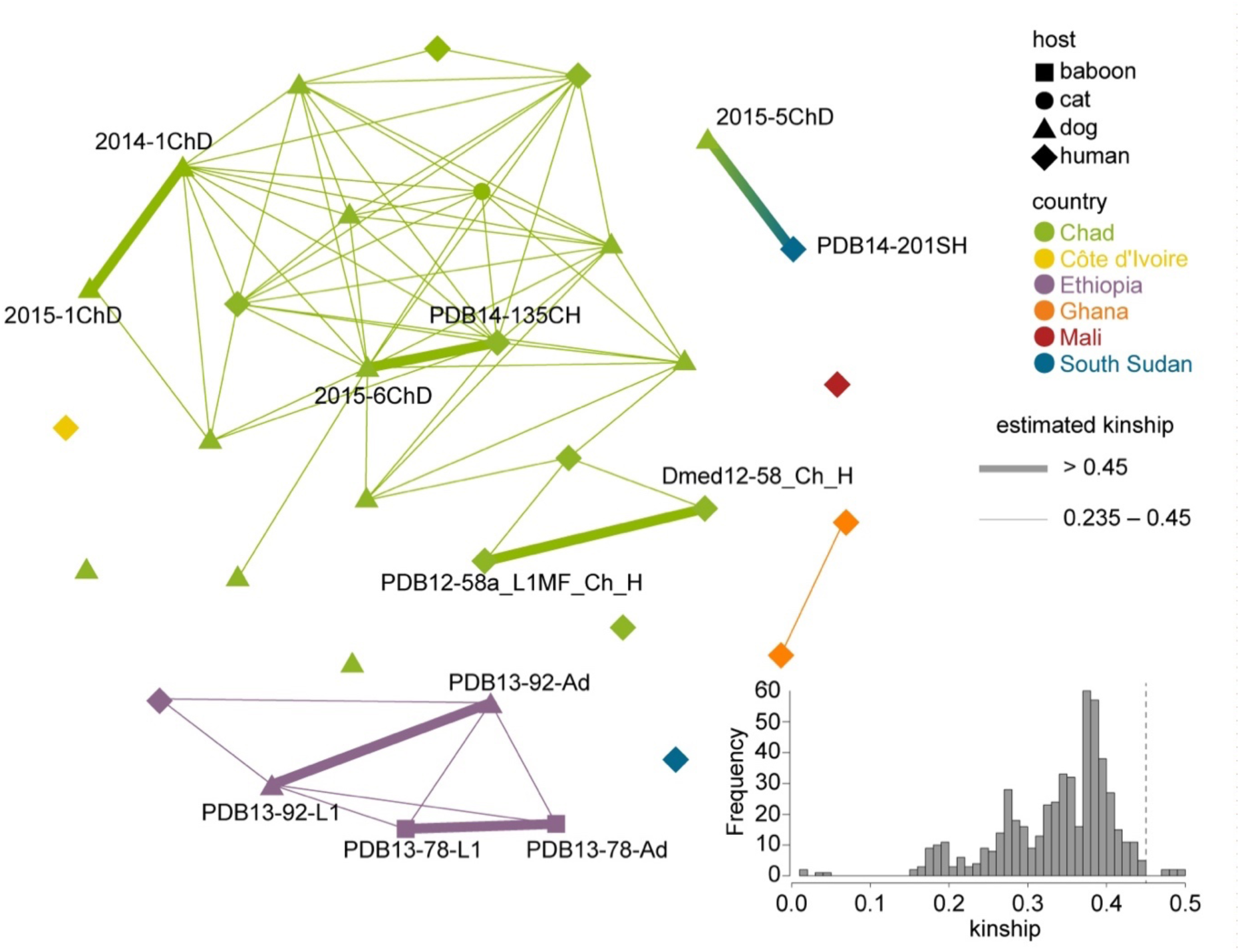
Relatedness between *Dracunculus medinensis* samples. Nodes on the graph represent worm samples, coloured by their country of origin, and node shapes indicate host species. Lines connect samples with high levels of identity by descent, indicative of direct relatedness. Thick lines indicate kinship > 0.45, whereas thinner lines indicate kinship between 0.45 and 0.235. For clarity, samples with other kinship coefficients are not connected in the graph, and sample names are shown only for those samples in high-relatedness pairs. Inset panel shows the distribution of kinship coefficients across all pairs of samples.

The three pairs we identify are all of significant epidemiological interest. One appears to confirm cross-border transmission, proposing that a worm emerging from a dog in Chad in 2015 was caused by a human case detected in South Sudan in 2014. A second pair links a human case in 2014 in Chad with a dog infection in 2015, apparently confirming transmission is possible between human cases and dog infections, while a third links two Chad dog infections in 2014 and 2015. One important note of caution is that all three of these events would imply long range transmission of the infection, with 1812km, 378km and 432km separating the three pairs of infections above, respectively; the two transmission events within Chad also imply movement in different directions on the Chari river basin (Figure 7).

**Figure 7.**
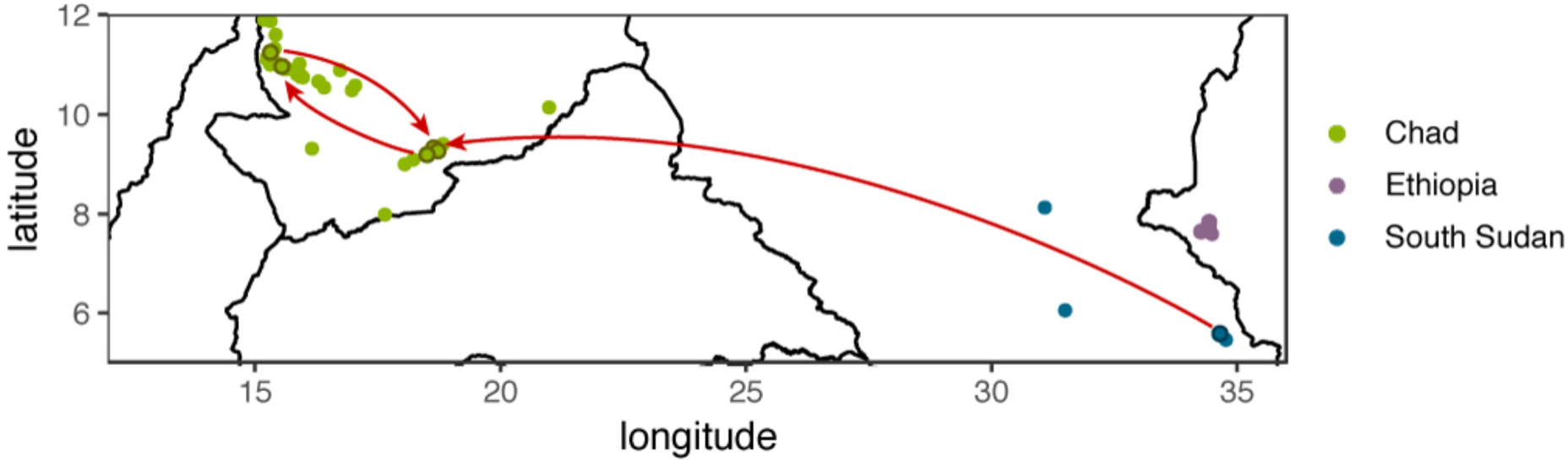
Transmission events implied by three parent-offspring pairs inferred from high genome-wide identity by descent between worms isolated in consecutive years. Sample locations are indicated by dots, colour-coded by country of isolation. Red arrows indicate inferred parent-offspring relationships between samples; samples involved in these links are highlighted by dark rings around the point at which the infection was detected. The locations of detection may not represent the locations at which infections were acquired, or the location of residence of the hosts.

## Discussion

Our data shows that a single population of *D. medinensis* is responsible for both dog infections and human cases in Chad, with genetic structure in *D. medinensis* being apparently driven by geographic separation rather than definitive host species. Our data suggest that all Guinea worm infections in in African mammals are caused by a single species, *D. medinensis.* The two other species of *Dracunculus* with mammalian hosts for which we have sequence data are highly divergent from any *D. medinensis* specimen we investigated. Genetic variation does exist within *D. medinensis* in Africa, but follows a spatial pattern, with populations from South Sudan and Ethiopia being more closely related to worms from Chad, and more divergent population of *D. medinensis* being present in West African countries prior to the recent elimination of the parasite from that region. The set of samples we have investigated from Chad and East Africa show a particularly high genetic diversity, but also strong signals of a recent population bottleneck, presumably driven by the ongoing work to eradicate Guinea worm in those areas.

We have identified three pairs of worms with high kinship that emerged in consecutive years, which we propose may represent transmission events. If so, our data confirm that cross-border transmission of Guinea worm infection can occur (from South Sudan to Chad in this case) and that infections can be passed from dog to dog and from humans to dogs. Unfortunately, we did not observe dog to human transmission directly in these data, although this is likely to be due to the small number of transmission events we could reconstruct, rather than because these transmissions are rare. Interpreting kinship in an inbred population is difficult, so these genealogical links must be considered only provisional, although we note that all three pairs of larval-adult samples for which we had good sequence coverage were correctly identified by this approach. While our data do not speak directly to the changes in lifecycle that might be driving transmission through dog hosts in Chad, both the long-range nature of these 3 transmission events and the fact that they imply different directions of movement along the Chari river basin would seem to lend some support to the idea that a paratenic or transport host could be involved. In particular, as one event is between two dog hosts, human movement may be less likely to be involved. It has been demonstrated experimentally that *D. medinensis* can pass through tadpoles as paratenic hosts and fish as transport hosts and that both routes can successfully infect ferrets (Cleveland et al., 2017; Eberhard, Yabsley, et al., 2016). Furthermore, a frog naturally infected with *D. medinensis* has been found in Chad (Eberhard, Cleveland, et al., 2016). Wildlife infections are also being reported, for example with a number of infections recently reported in Baboons in Ethiopia (Hopkins et al., 2018).

Our coalescent models of the Guinea worm population genetic data appear to confirm the geographical structure of these populations, and that worms from human cases and dog infections in Chad form a single population. The estimates of population divergence dates imply that the genetic structure we observe between different regions of Africa predates recent control efforts and likely represents historical population structure. The oldest subdivision we observe, at around 20,000 years ago, coincides with the last glacial maximum when Africa was likely to be extremely arid, even compared to present-day conditions (Hoag & Svenning, 2017). Similarly the more recent divergence between East African and Chad populations at around 4,000 years ago is during the drying-out of the Sahara at the end of the African humid period (Hoag & Svenning, 2017), which was probably accompanied by a major collapse in human habitation of much of this region (Manning & Timpson, 2014). Although our qualitative results appear robust, there are more caveats with the specific quantitative results. In particular, these estimates depend on assumptions about the mutation rate and generation time of *D. medinensis*. It is generally accepted that Guinea worm infections take approximately 10-14 months to reach patency in human infections (Cairncross et al., 2002; Muller, 1971). Less certain is whether larvae can remain viable in copepods or within a paratenic host for extended periods of time. No direct measurement of the mutation rate is available for *D. medinensis* or any related parasitic nematode, and while mutation rates are reasonably consistent across eukaryotes with similar genome sizes (Lynch, 2010), variation of several fold from the value we have assumed would not be very surprising. We also note that the relative values of divergence time and population size estimates will remain unchanged under different mutation rates.

Our quantitative model suggests that all three present-day populations have large average effective population size (N_e_) (of the order of thousands to low tens of thousands) over thousands of years. The modelling approach we have used is not able to detect more recent changes in these populations, and interpreting these estimates is challenging, as genetic effective population sizes are influenced by many factors such as breeding systems, demography and selection. In particular, historical fluctuations in population size have a strong influence on N_e_, approximated by the geometric mean of the population sizes across generations (Charlesworth & Charlesworth, 2010 pp225-226). The high N_e_ in Chad appears to exclude the possibility that the population of worms in Chad either disappeared or was reduced to a very small bottleneck during the decade without reported human cases; it is difficult to reconcile with a population size during this time much below hundreds of worms. In the absence of Chad samples prior to 2000, or more extensive sampling from neighbouring countries we cannot exclude the possibility that the Chad worms we analyse – which all emerged in Chad following the 10-year gap in reported cases – migrated from elsewhere.

However, we see few Chad worms that are closely related to worms from any of the neighbouring countries for which we had access to samples, so this possibility is purely speculation, and it would seem that quite large-scale influx would be required to explain the level of diversity we see in Chad by migration. Without historical samples, it also remains uncertain to what extent the population structure we see in African Guinea worm today would have been different 30 years ago, when the census population size of the worms was more than three orders of magnitude higher and worms were still widespread in Africa. Our coalescence model suggests that at least Chad and the East African populations we have sampled were still largely distinct at this time, but we have not been able to obtain worm samples suitable for molecular analysis from much of the ancestral range of *D. medinensis*.

A limitation is the nature of samples available to us, and in particular the very small quantity of genuinely adult material present in specimens despite these being very large for a parasitic nematode. Enrichment methods targeting parasite over host DNA cannot enrich for adult versus larval DNA and it is operationally difficult to alter the way that material is collected in the field in the context of the eradication campaign. The nature of our existing samples as mixtures of many diploid individuals makes some forms of analysis challenging: for example, many of the most sensitive signatures of inbreeding we expect to see appearing as the population size declines rely on changes in the level and distribution of homozygous and heterozygous sites (Diez-Del-Molino, Sanchez-Barreiro, Barnes, Gilbert, & Dalen, 2018). These are not readily apparent in analysis of the data presented here, presumably because of the mixture of genotypes present in each sample. These may be particularly complex if, like many other parasitic nematodes *D. medinensis* is polyandrous (Redman et al., 2008; Zhou, Yuan, Tang, Hu, & Peng, 2011; Doyle et al., 2018). We are currently generating sequence data from individual L1 larvae which should let us look for these signals, dissect the contribution of different males to a brood, and infer recombination and mutation rates in *D. medinensis*, avoiding the need to rely on estimates from *C. elegans*, which is both very distantly related to *D. medinensis* and has, of course, a very different life history. We have recently demonstrated the feasibility of this approach in a different parasitic nematode system (Doyle et al., 2018). Efforts to extract useful genome-wide information from the low-quality *D. medinensis* samples not analysed here are ongoing, with results from a sequence capture approach showing some promise. Our results are consistent with the findings of previously published targeted genotyping with mitochondrial and microsatellite markers, which also produced additional insights into the population genetics of *D. medinensis* from a much more extensive set of parasite samples (Thiele et al., 2018).

Finally, the data we present here, together with other data from Guinea worm populations (Bimi et al., 2005; Eberhard et al., 2014; Thiele et al., 2018) preserve something of the genetics of *D. medinensis* in the final foci of infection. The genome sequence should help preserve some of the biology of this important human pathogen following the extinction of *D. medinensis* with eradication, but more importantly we expect these data to be crucial in the final steps aftermath of the eradication process. By defining much of the currently existing diversity of Guinea worm, these data will act as a reference to determine whether future cases for which the source of infection is unclear represent continuing transmission from these foci or previously unidentified worm populations. The emergence of large numbers of dog infections in Chad could not have been predicted, and the eradication campaign could uncover other unexpected aspects of Guinea worm biology or epidemiology. For example, a recent surprise is the emergence of Guinea worm infections in Angola, which has no previous history of Guinea worm disease (Centers for Disease Control and Prevention, 2018). It is likely that reports of emerging worms will appear post-eradication (Mbong et al., 2015): given the paucity of morphological features defining *D. medinensis*, molecular tools will be key in providing certainty about the pathogen involved, and thus ultimately in allowing the WHO to declare that the world is free of Guinea worm.

Our work has clear implications for other parasite systems as we move into an era intended to see enhanced control efforts, regional elimination and even eradication for several neglected tropical disease parasites (World Health Organisation, 2012). The Guinea worm eradication program has in many ways set the scene for these efforts in other parasites. The small size of the remaining Guinea worm populations means it should be particularly feasible to employ whole-genome approaches to track changes in Guinea worm populations during the final stages of eradication (Cotton, Berriman, Dalen, & Barnes, 2018), but the particular difficulties in generating high-quality sequence data and in interpreting these data for *D. medinensis* highlight the fact that every pathogen system is unique, and genetic surveillance will likely face unique challenges in each case. Whether the particular challenges of an apparently emerging zoonotic transmission cycle in the endgame of eradication are unique remains to be seen as programs for other pathogens advance. It seems clear that the endgame of elimination has different requirements to much of the process of reducing disease burden (Klepac, Metcalf, McLean, & Hampson, 2013) and the strong selection pressure on pathogen populations to evade control measures near eradication will result in evolutionary responses. The ecological changes apparently occurring in Guinea worm may be the equivalent of the evolution of drug resistance in chemotherapy-lead campaigns (Whitty, 2014).

## Conclusion

Our results are entirely consistent with a single population of *D. medinensis* infecting both dogs and humans in Chad. We show genetic variation within *D. medinensis* is largely geographical, with significant differentiation between populations present in Chad, and those present in countries in East Africa (South Sudan and Ethiopia) and West Africa (Côte d’Ivoire, Ghana, Mali and Niger). Worms that were genetically very similar were recovered from human cases and animal infections in both Chad and Ethiopia. We find a particularly diverse population of worms in Chad and East Africa that appears to be shrinking, presumably due to the eradication program. Coalescent models confirm that a single population of worms infects both dogs and humans in Chad, and the long-term effective population size suggests that a significant Guinea worm population persisted in Chad during the ten-year period prior to 2010 during which no cases were reported. Kinship analysis shows that the Guinea worm population is highly inbred, as we might expect in a small and shrinking population, and suggests direct relatedness between 3 pairs of worms, including two recovered from human cases in one year and recovered from dogs in a subsequent season. In the context of epidemiological data and previous genetic data, this suggests that dog infections are likely to be central to maintaining Guinea worm transmission in Chad. Continued efforts to understand the biology of transmission in Chad, as well as sustained surveillance among both human and non-human hosts, will help ensure the continuing success of the eradication program.

## Methods

Worm material from *D. medinensis* was collected by the national Guinea worm eradication programs in the relevant countries, except that material from experimentally infected ferrets were obtained as previously described (Eberhard, Yabsley, et al., 2016). *D. insignis* material was collected from an American mink (*Neovison vison*) and *D. lutrae* material was collected from an otter (*Lutra canadensis*) in Ontario, Canada (Elsasser et al., 2009).

Genomic DNA was extracted from either 5-15mm sections of adult female worm specimens or from the pool of L1 larvae visible in sample tubes, wherever larvae were visible. DNA extraction was perfomed using the Promega Wizard kit, but with worm specimens cut into small pieces before digestion with 200µg of Proteinase K overnight in 300 µl of lysis buffer, then following the protocol described in the manual. PCR-free 200 – 400 bp paired-end Illumina libraries were prepared from genomic DNA as previously described (Kozarewa et al., 2009) except that Agencourt AMPure XP beads were used for sample clean up and size selection. DNA was precipitated onto the beads after each enzymatic stage with a 20% (w/v) Polyethylene Glycol 6000 and 2.5 M sodium chloride solution, and beads were not separated from the sample throughout the process until after the adapter ligation stage. Fresh beads were then used for size selection. Where there was insufficient DNA for PCR-free libraries, adapter-ligated material was subjected to ∼8-14 PCR cycles. Libraries were run on an Illumina platform (HiSeq 2000, 2500 or HiSeq X) to generate 100 base pair or 150 base pair paired-end reads.

Sequence data was compared to a reference genome assembled from a worm collected in Ghana in 2001. The sequence data and automated assembly of v2.0 of this reference is described fully elsewhere (International Helminth Genomes Consortium, 2017). The v3.0 reference used here has undergone some manual improvement, with REAPR (Hunt et al., 2013) used to identify problematic regions of the assembly to be broken, followed by iterative rounds of re-scaffolding as indicated by read-pair and coverage information visualised in GAP5 (Bonfield & Whitwham, 2010) and automated gap-filling with IMAGE (Tsai, Otto, & Berriman, 2010) and Gap-filler v1.11 (Nadalin, Vezzi, & Policriti, 2012) and a final round of sequence correction with iCORN v2.0 (Otto, Sanders, Berriman, & Newbold, 2010). Assembly statistics for v2.0 and v3.0 of the *D. medinensis* genome are shown in Supplementary Table 1.

Mapping was performed with SMALT v.0.7.4 (http://www.sanger.ac.uk/science/tools/smalt-0), mapping reads with at least 95% identity to the reference, mapping paired reads independently and marking them as properly paired if the reads within a pair mapped in the correct relative orientation and within 1000bp of each other (parameters –x –y 0.95 –r 1 –i 1000). To avoid problems with mitochondrial data, mapping was also performed similarly against a reference containing mitochondrial genomes for dog, human and ferret. Duplicate reads were removed using Picard v2.6 MarkDuplicates. The BAM files produced were used as input to Genome Analysis Toolkit v3.4.0 for variant calling, following the ‘best practice’ guidelines for that software release: briefly, reads were realigned around indel sites, after which SNP variants were called using HaplotypeCaller with ploidy 2. Variants were then removed where they intersected with a mask file generated with the GEM library mappability tool (Derrien et al., 2012) with kmer length 100 and 5 mismatches allowed, or were within 100bp of any gap within scaffolds. Finally, SNPs were then filtered to keep those with DP >= 10; DP <= 1.75*(contig median read depth); FS <= 13.0 or missing; SOR <= 3.0 or missing, ReadPosRankSum <= 3.1 AND ReadPosRankSum >= −3.1; BaseQRankSum <= 3.1 AND BaseQRankSum >= −3.1; MQRankSum <= 3.1 AND MQRankSum >= −3.1; ClippingRankSum <= 3.1 AND ClippingRankSum >= −3.1. An additional mask was applied, based on the all-sites base quality information output by GATK HaplotypeCaller. The filters applied were DP >= 10, DP <= 1.75*(contig median read depth) and GQ >= 10. Finally, sites with only reference or missing genotypes were then removed. Variant calling on the mitochondrion was performed similarly, except reads were first filtered to retain only those for which both reads in pair mapped uniquely to the mitochondrion in the correct orientation with mapping quality at least 20, the read depth filter was 10 for all samples, all heterozygous calls were removed and the mask file was generated manually by examining dot plots and removing regions with a high density of heterozygotes.

Synteny between the *D. medinensis* v3.0 assembly and the published *O. volvulus* v4.0 assembly was confirmed using promer from the Mummer package (Kurtz et al., 2004) to identify regions of >50% identity between the two sequences over 250 codons. These results were then visualised using Circos v0.67pre5 (Krzywinski et al., 2009). The phylogenetic tree for *D. medinensis* samples was based on the proportion of alleles matching between each pair of samples at those sites for which both samples in a pair had a genotype call that passed the filter criteria. A phylogeny based on these distances was inferred by neighbour-joining using the program Neighbour from Phylip v3.6 (Felsenstein, 2005). Principal components analysis was performed on a matrix of genotypes for sites with no missing data in R v3.3.0 (R Core Team, 2015) using the prcomp command. Population genetic summary statistics within and between populations were calculated for 10kb window of SNPs containing between 5 and 500 variants, using ANGSD v0.919-20-gb988fab (Korneliussen, Albrechtsen, & Nielsen, 2014). This software estimates neutrality tests (Korneliussen, Moltke, Albrechtsen, & Nielsen, 2013) or genetic differentiation between populations (Fumagalli et al., 2013) following a probabilistic framework that employs genotype likelihoods. It is intended to be more robust to genotyping error than traditional calculations using the genotypes directly. Only sites with a minimal depth of 5 reads, a minimal base and mapping quality Phred score of 30 and a call rate of at least five individuals were used, and genotype likelihoods were estimated under the samtools (Li et al., 2009) framework (GL = 1). In the absence of known ancestral states, folded site frequency spectra were generated to derive nucleotide diversity π, Watterson’s θ and Tajima’s D. F_ST_ estimates were computed from maximum-likelihood joint site frequency spectra between pairs of populations derived using the reference genome as the ancestral state. Estimates were generated for 10-Kb sliding windows (with 1 Kb overlaps) containing between 5 and 500 variants. We report genome-wide averages across these windows, and confidence intervals for these statistics were calculated for 10kb from 100 bootstrap replicates, resampling from 10kb windows. Unless otherwise specified, plots were produced in R v3.3.0 with ggplot2 (Wickham, 2009).

Bayesian clustering was performed with MavericK v1.0 using thermodynamic integration to estimate the number of clusters (*K*) best describing the data (Verity & Nichols, 2016). MavericK was run for 3 independent runs of 1,000 burnin generations and then 10,000 generations for inference, and with each rung of the thermodynamic integration run for 1,000 burnin generations and 5,000 generations for inference, for the default 21 rungs. For comparison, Structure v2.3.4 was run (Pritchard, Stephens, & Donnelly, 2000), using the deltaK method to select a value of *K*. Input to both of these was a set of 19,983 SNP variants samples across the *D. medinensis* scaffolds at 5kb intervals. Structure was run using an admixture model, with a burn-in of 100,000 generations and using another 100,000 generations for inference.

Population history of *D. medinensis* was inferred using BPP v4 to infer the number and branching pattern of populations, and then GPhoCS v1.2.2 (Gronau, Hubisz, Gulko, Danko, & Siepel, 2011) to infer branching times and effective population sizes on the maximum posterior probability history. GPhoCS requires populations and the phylogeny to be specified a-priori, but is able to perform inference using a larger set of loci more efficiently. For GPhoCS a total of 781 loci were chosen as contiguous 1kb regions spaced every 100kb across all autosomal scaffolds. Results were scaled to time and effective population time using a mutation rate of 2.7 × 10^−9^ per generation, as estimated for *Caenorhabditis elegans* (Denver et al., 2009) and a generation time of 12 months: *D. medinensis* females emerge 10-14 months after infection (Muller, 1979). At least three (3-6) independent MCMC chains were run for each of 5 different prior assumptions, with each chain running for at least 25,000 MCMC generations. In each case, identical priors were used for all θ and τ (population sizes and divergence times, respectively) parameters; priors for GPhoCS are specified as gamma distributions parameterised with a shape (α) and rate (β) parameters (hyperparameters).

We held the β hyperparameter constant at 0.1, and chose α values varying by 4 orders of magnitude, from 10^−4^ to 10^−8^, so that the means of the prior distributions varied from 6.43335-64,335 for θs following scaling and from 25.7342 to 257,342 for τ parameters. The variance of the prior distributions also varied linearly with changes in the α hyperparameter. Convergence was confirmed by visual inspection of the chains for each prior. For inference, the first 15,000 generations of each chain were removed and the remaining steps concatenated; highest posterior density estimates and effective sample sizes were calculated using the R packages HDInterval and mcmcse respectively. For all parameters the effective sample size was at least 250.

BPP attempts to identify reproductively isolated populations and estimate the phylogeny underlying those populations in a joint Bayesian framework (Yang & Rannala, 2010; Rannala & Yang, 2017). Population size parameters were assigned the default inverse gamma priors with mean 0.002 and shape parameter (alpha)=3, the root divergence time an inverse gamma prior with mean 0.001 and alpha 3, other divergence time parameters default Dirichlet prior. Each analysis is run at least twice to confirm consistency between runs, and each chain was run for 10,000 burnin generations and 50,000 generations for inference. Convergence was assessed by inspection of these chains in Tracer v.1.6. For BPP, a subset of 100 loci was chosen at random from these 781 loci. Three different random sets of loci gave essentially identical results (97.2%, 98.2% and 98.4% support for the same maximum-probability reconstruction; duplicate runs of the same loci varied by less than 0.5%).

Kinship between samples was calculated using King v1.4 (Manichaikul et al., 2010). Distances between latitude and longitude points were calculated using the online calculator at the US National Oceanic and Atmospheric Administration at https://www.nhc.noaa.gov/gccalc.shtml.

## Declarations

### Ethics approval and consent to participate

Specific ethical approval was not required as material was derived from standard containment and treatment procedures sanctioned by WHO and national governments and performed by national control program staff, and molecular testing is part of the standard case confirmation procedure. Human case samples were anonymized prior to inclusion in this study.

### Consent for publication

Not applicable

### Availability of data and materials

All data generated in this study are available from the European Nucleotide Archive short read archive, a project ERP117282; accession numbers for individual samples are shown in supplementary table 1.

### Authors’ contributions

CD, EAT, SRD, GS, AT and JAC analysed data. GS, ML, HMB, TH and ZA performed molecular biology. OT, MW, MSYL, COC, AW, AIS-H, JF, CAC, MJY, ER-T and MLE provided material. NH co-ordinated the generation of sequence data. ER-T, MB, MLE and JAC designed the study. JAC wrote the manuscript draft with contributions from CD and EAT. NH, GS, SRD, CAC, MJY, ER-T, MB and MLE also reviewed and edited the manuscript. All authors read and approved the final manuscript.

## Acknowledgements

We thank the national Guinea worm eradication programs for collecting and making worm specimens and associated data available and the support of the Guinea worm program teams from the Carter Center and Centers for Disease Control and Eradication.

---

**Supplementary Figure 1.**
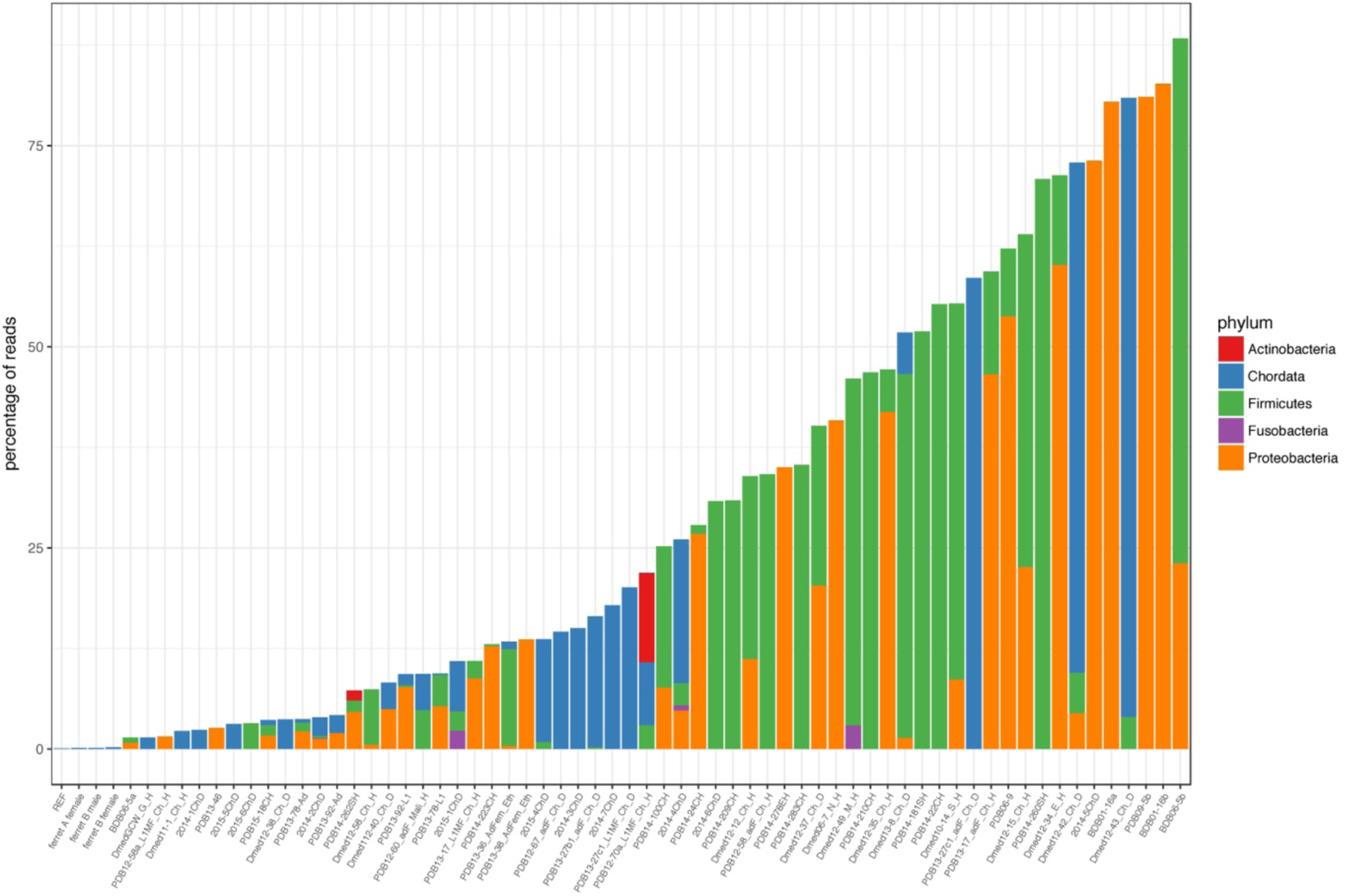
Sources of contamination in sequencing libraries. Number of reads inferred by k-mer analysis to originate from different phyla. Data are shown for all phyla to which at least 50,000 reads were assigned. 68 samples are shown: those not shown here did not match any phyla with his cut-off. Note that no nematode sequences are in the database used for this search (see methods).

**Supplementary Figure 2.**
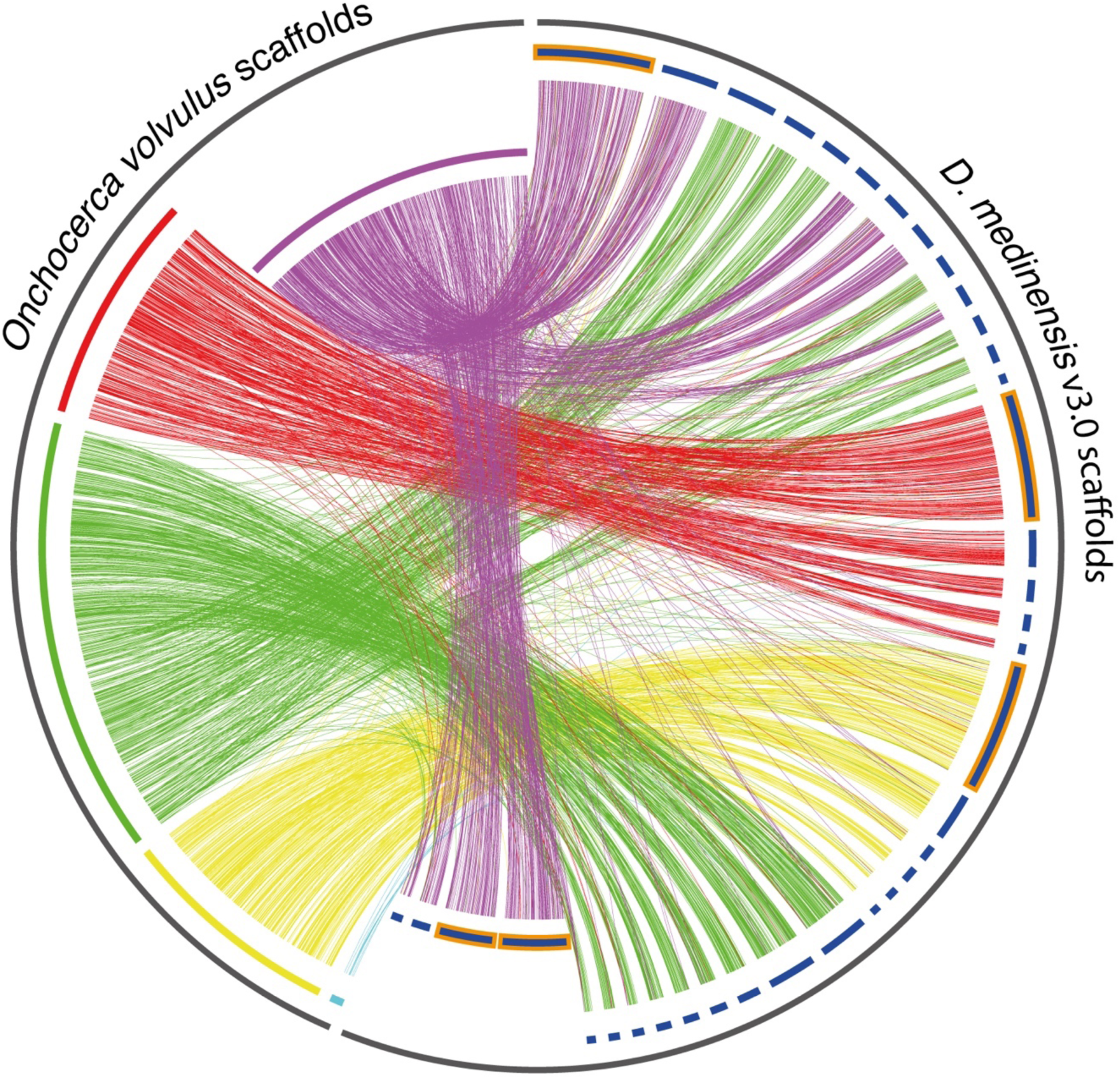
Synteny between *D. medinensis* and *O. volvulus* scaffolds. Lines connect sequences for which the conceptual amino acid translations are at least 50% identical over 250 amino acids. *D. medinensis* scaffolds highlighted in orange are those shown in Figure 2a. Note that one of the longest scaffolds matches to the opposite end of the X-chromosome scaffold in *O. volvulus* to the scaffolds with reduced coverage in male worms. This region of *O. volvulus* X was not part of the ancestral filarial X chromosome (Cotton et al., 2016), and so is not expected to be part of *D. medinensis* X and is thus labelled as autosomal in Figure 2a, and considered as autosomal in our analyses here. *D. medinensis* scaffolds with reduced male coverage are shown inset.

**Supplementary Figure 3.**
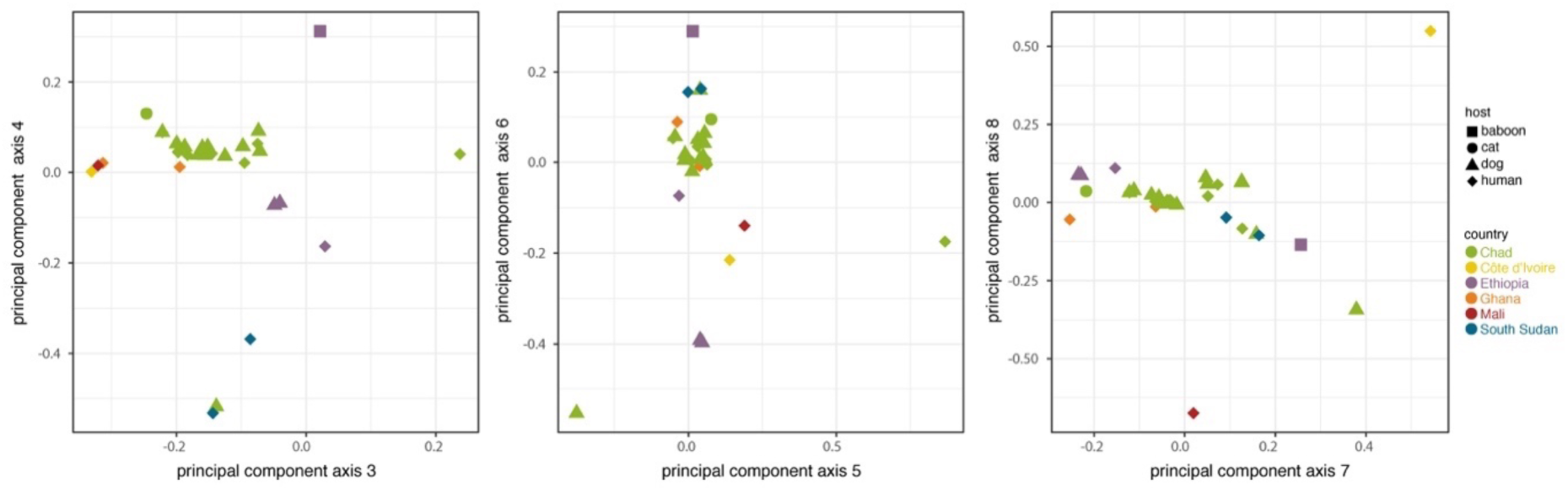
PCA axes 3-8 for *Dracunculus medinensis* variation data; axes 1 and 2 are shown in main text Figure 3c.

**Supplementary Figure 4.**
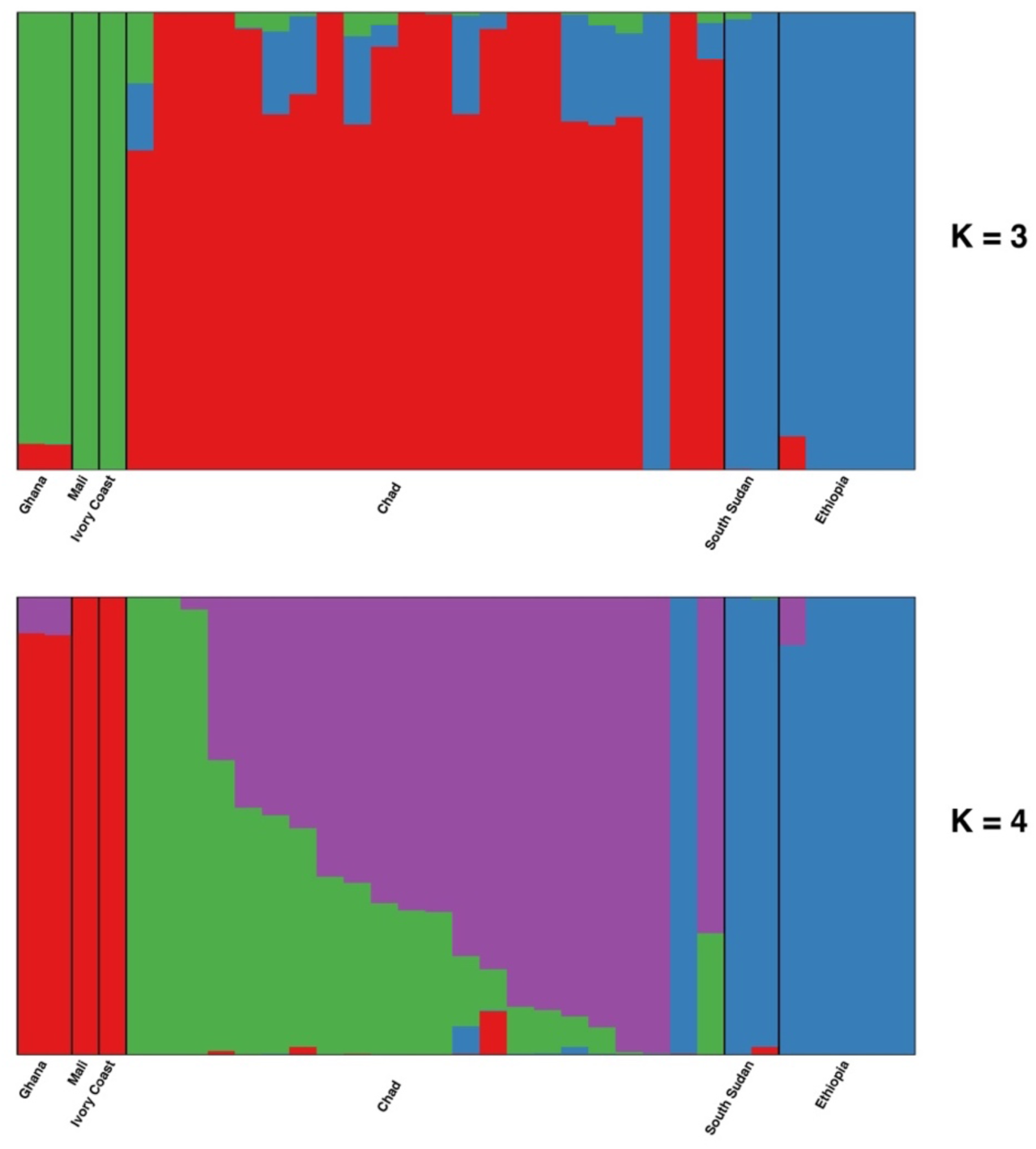
Bayesian assignment of individual samples to populations, for k=3 and k=4 hypothetical populations. Vertical bars represent individuals, with the proportion of each color in each bar representing the proportion of inferred ancestry of that individual from the population. Note that the order of individual samples within each country differs between the two panels.

**Supplementary Figure 5.**
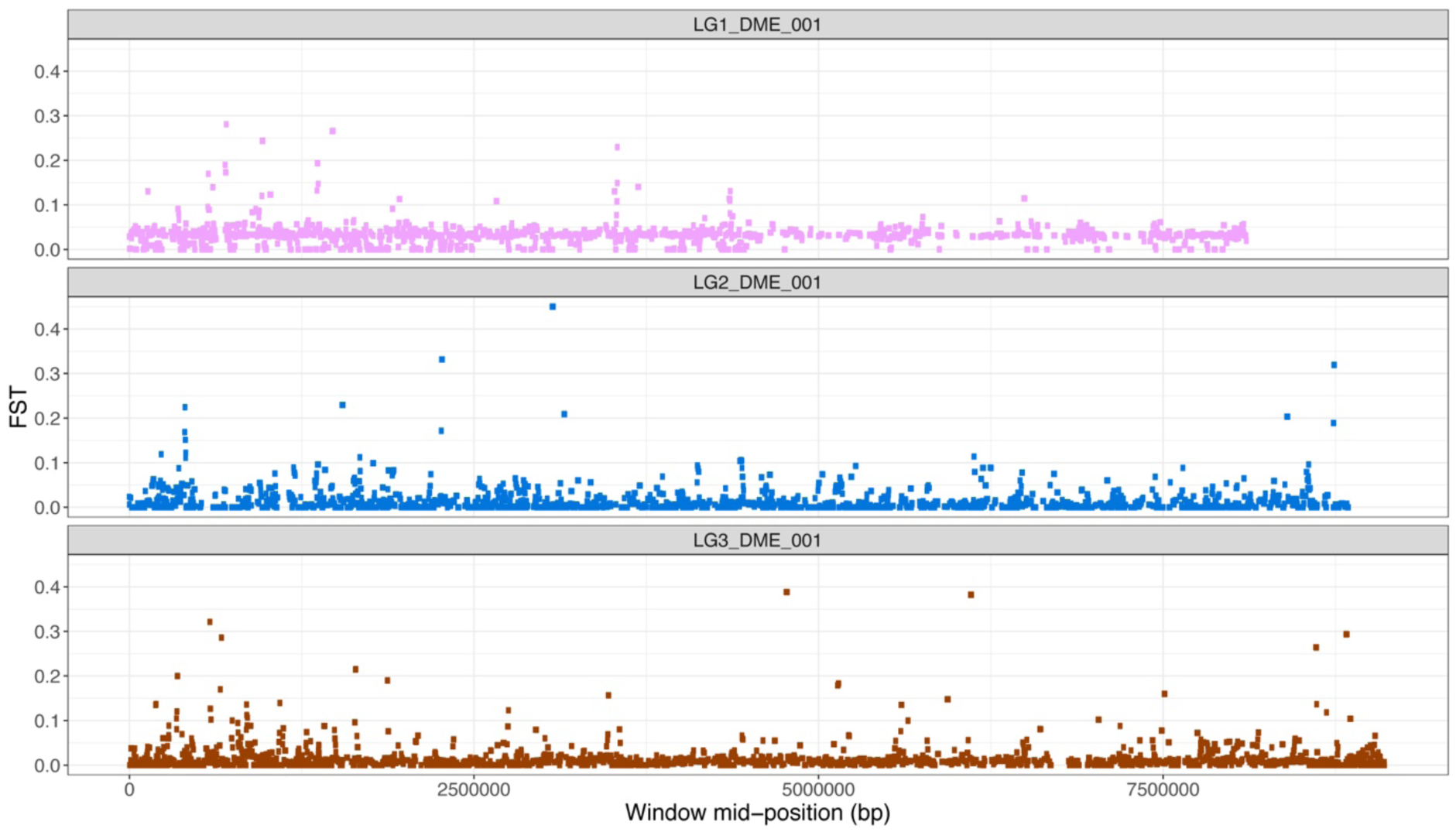
F_st_ between *Dracunculus medinensis* samples from dogs and humans in Chad, across the three longest autosomal scaffolds. Values shown are mean F_st_ for non-overlapping 1kb windows centered at the position shown on the x axis.

**Supplementary Table 1.** Details of *Dracunculus medinensis* samples, sequencing data and sequencing libraries used in this study. Note that mean and median coverage are defined over the whole nuclear genome assembly for both MIT and NUC samples. Reads and mapping statistics are for the sum across all sequenced libraries and lanes. ENA= European Nucleotide Archive.

| sample name | country | host | variant data | total reads | reads mapping | percent mapping | mean coverage | median coverage | number of lanes | ENA accession numbers |
| --- | --- | --- | --- | --- | --- | --- | --- | --- | --- | --- |
| PDB14-138CH | Chad | Human | no library made |  |  |  |  |  |  |  |
| PDB14-23CH | Chad | Human | no library made |  |  |  |  |  |  |  |
| PDB14-25CH | Chad | Human | no library made |  |  |  |  |  |  |  |
| PDB14-287MH | Mali | Human | no library made |  |  |  |  |  |  |  |
| BDB06-5b | Cote d'Ivoire | Human | MIT | 2,229,520 | 21,157 | 0.95 | 0.03 | 0 | 1 | ERR460382 |
| PDB06-9 | Cote d'Ivoire | Human | MIT | 2,115,778 | 365,211 | 17.26 | 0.53 | 0 | 1 | ERR460383 |
| PDB14-181SH | South Sudan | Human | NONE | 228,914 | 64,654 | 28.24 | 0.09 | 0 | 1 | ERR1081345 |
| PDB14-260SH | South Sudan | Human | NONE | 1,095,228 | 2,610 | 0.24 | 0.00 | 0 | 2 | ERR1081343 |
| PDB14-262SH | South Sudan | Human | NONE | 1,007,068 | 2,189 | 0.22 | 0.00 | 0 | 2 | ERR1081342,ERR1730377 |
| PDB14-201SH | South Sudan | Human | NUC | 11,201,696 | 9,920,172 | 88.56 | 12.05 | 10 | 2 | ERR1081344,ERR1243214 |
| Dmed10-14_S_H | South Sudan | Human | NUC | 138,443,686 | 24,876,967 | 17.97 | 23.95 | 19 | 4 | ERR273907,ERR273929,ERR563493,ERR563499 |
| 2014-1ChD | Chad | Dog | NUC | 25,387,772 | 15,168,391 | 59.75 | 18.49 | 14 | 2 | ERR1081328,ERR1243211 |
| 2014-2ChD | Chad | Dog | NUC | 118,848,030 | 59,278,247 | 49.88 | 71.60 | 61 | 2 | ERR1081357,ERR1243221 |
| 2014-3ChD | Chad | Dog | NUC | 20,821,588 | 13,348,157 | 64.11 | 16.12 | 14 | 2 | ERR1081358,ERR1243222 |
| 2014-4ChD | Chad | Dog | NUC | 34,795,436 | 17,975,620 | 51.66 | 21.76 | 19 | 2 | ERR1081359,ERR1243223 |
| 2014-5ChD | Chad | Dog | MIT | 429,444 | 55,993 | 13.04 | 0.08 | 0 | 1 | ERR1081346 |
| 2014-6ChD | Chad | Dog | MIT | 680,944 | 203,121 | 29.83 | 0.29 | 0 | 1 | ERR1081347 |
| 2014-7ChD | Chad | Dog | MIT | 665,416 | 216,817 | 32.58 | 0.31 | 0 | 1 | ERR1081348 |
| 2014-8ChD | Chad | Dog | NONE | 338,870 | 40,552 | 11.97 | 0.06 | 0 | 1 | ERR1081349 |
| 2015-1ChD | Chad | Dog | NUC | 24,776,440 | 14,464,333 | 58.38 | 17.47 | 13 | 2 | ERR1081350,ERR1243215 |
| 2015-2ChD | Chad | Dog | MIT | 8,900,594 | 7,848,792 | 88.18 | 9.51 | 8 | 2 | ERR1081351,ERR1243216 |
| **2015-3ChD** | Chad | Dog | NUC | 11,763,796 | 10,321,366 | 87.74 | 12.54 | 11 | 2 | ERR1081352,ERR1243217 |
| **2015-4ChD** | Chad | Dog | NUC | 33,738,410 | 10,758,127 | 31.89 | 13.03 | 11 | 2 | ERR1081326,ERR1243209 |
| **2015-5ChD** | Chad | Dog | NUC | 11,309,548 | 9,933,132 | 87.83 | 12.06 | 10 | 2 | ERR1081327,ERR1243210 |
| **2015-6ChD** | Chad | Dog | NUC | 17,397,832 | 11,787,265 | 67.75 | 14.24 | 13 | 2 | ERR1081353,ERR1243218 |
| **2015-7ChD** | Chad | Dog | MIT | 324,328 | 198,454 | 61.19 | 0.29 | 0 | 1 | ERR1081354 |
| **2015-8ChD** | Chad | Dog | MIT | 8,057,292 | 6,838,667 | 84.88 | 8.27 | 7 | 2 | ERR1081355,ERR1243219 |
| **BDB06-5a** | Cote d'Ivoire | Human | NUC | 89,024,662 | 67,376,292 | 75.68 | 65.70 | 55 | 5 | ERR460381,ERR563547,ERR563553,ERR563559,ERR563564 |
| **BDB01-16a** | Togo | Human | NONE | 1,800,622 | 1,522 | 0.08 | 0.00 | 0 | 1 | ERR460379 |
| **BDB01-16b** | Togo | Human | NONE | 1,798,440 | 1,682 | 0.09 | 0.00 | 0 | 1 | ERR460380 |
| **Dmed06-7_N_H** | Niger | Human | NONE | 9,731,926 | 25,777 | 0.26 | 0.02 | 0 | 2 | ERR273906,ERR273928 |
| **Dmed11-1_Ch_H** | Chad | Human | NUC | 60,468,586 | 50,606,117 | 83.69 | 48.74 | 39 | 6 | ERR273901,ERR273923,ERR563491,ERR563497,ERR563525,ERR563536 |
| **Dmed112-40_Ch_D** | Chad | Dog | MIT | 16,826,492 | 535,336 | 3.18 | 0.52 | 0 | 2 | ERR273914,ERR273936 |
| **Dmed12-12_Ch_H** | Chad | Human | NONE | 4,146,036 | 15,934 | 0.38 | 0.02 | 0 | 2 | ERR273902,ERR273924 |
| **Dmed12-15_Ch_H** | Chad | Human | NONE | 4,465,058 | 3,230 | 0.07 | 0.00 | 0 | 2 | ERR273903,ERR273925 |
| **Dmed12-34_E_H** | Ethiopia | Human | MIT | 12,117,156 | 1,727,610 | 14.26 | 1.66 | 1 | 2 | ERR273908,ERR273930 |
| **Dmed12-35_Ch_H** | Chad | Human | NONE | 11,603,600 | 313,203 | 2.7 | 0.30 | 0 | 2 | ERR273904,ERR273926 |
| **Dmed12-37_Ch_D** | Chad | Dog | MIT | 14,276,312 | 55,830 | 0.39 | 0.05 | 0 | 4 | ERR273912,ERR273934,ERR319496,ERR319497 |
| **Dmed12-38_Ch_D** | Chad | Dog | NUC | 69,686,582 | 61,063,480 | 87.63 | 58.83 | 52 | 6 | ERR273913,ERR273935,ERR563495,ERR563501,ERR563529,ERR563540 |
| **Dmed12-42_Ch_D** | Chad | Dog | MIT | 15,437,512 | 527,224 | 3.42 | 0.51 | 0 | 2 | ERR273915,ERR273937 |
| **Dmed12-43_Ch_D** | Chad | Dog | MIT | 15,124,806 | 1,363,831 | 9.02 | 1.31 | 0 | 2 | ERR273916,ERR273938 |
| **Dmed12-49_M_H** | Mali | Human | MIT | 11,720,346 | 21,964 | 0.19 | 0.02 | 0 | 2 | ERR273909,ERR273931 |
| **Dmed12-58_Ch_H** | Chad | Human | NUC | 255,743,466 | 216,521,130 | 84.66 | 208.59 | 180 | 8 | ERR273905,ERR273927,ERR563492,ERR563498,ERR563526,ERR563537,ERR563560,ERR563565 |
| **Dmed12-60_M_H** | Mali | Human | MIT | 1,886,708 | 1,437,002 | 76.16 | 1.38 | 1 | 2 | ERR273910,ERR273932 |
| **Dmed13-8_Ch_D** | Chad | Dog | MIT | 193,521,152 | 31,504,467 | 16.28 | 30.60 | 10 | 3 | ERR349721,ERR563503,ERR563514 |
| **DmedGCW_G_H** | Ghana | Human | NUC | 51,467,850 | 44,327,136 | 86.13 | 42.71 | 37 | 6 | ERR273911,ERR273933,ERR563494,ERR563500,ERR563528,ERR563539 |
| **PDB09-5a** | Niger | Human | MIT | 952,294 | 529,410 | 55.59 | 0.76 | 0 | 1 | ERR460384 |
| **PDB09-5b** | Niger | Human | NONE | 1,654,502 | 2,611 | 0.16 | 0.00 | 0 | 1 | ERR460385 |
| PDB12-58_adF_Ch_H | Chad | Human | MIT | 1,855,966 | 1,017,293 | 54.81 | 1.47 | 1 | 1 | ERR349722 |
| PDB12-58a_L1MF_Ch_H | Chad | Human | NUC | 78,014,152 | 66,426,989 | 85.15 | 64.98 | 58 | 5 | ERR349723,ERR563504,ERR563515,ERR563530,ERR563541 |
| PDB12-60_adF_Mali_H | Mali | Human | NUC | 43,539,932 | 28,816,385 | 66.18 | 27.92 | 23 | 5 | ERR349724,ERR563505,ERR563516,ERR563531,ERR563542 |
| PDB12-67_adF_Ch_D | Chad | Dog | NUC | 139,667,262 | 100,037,721 | 71.63 | 97.62 | 85 | 5 | ERR349729,ERR563508,ERR563519,ERR563532,ERR563543 |
| PDB12-67_L1MF_Ch_D | Chad | Dog | MIT | 819,306 | 634,946 | 77.5 | 0.92 | 1 | 1 | ERR349730 |
| PDB12-70a_L1MF_Ch_H | Chad | Human | NUC | 92,134,304 | 23,316,394 | 25.31 | 22.64 | 14 | 3 | ERR349726,ERR563506,ERR563517 |
| PDB12-70b_adF_Ch_H | Chad | Human | NONE | 402,610 | 187,091 | 46.47 | 0.27 | 0 | 1 | ERR349725 |
| PDB13-17_adF_Ch_H | Chad | Human | MIT | 1,510,120 | 432,055 | 28.61 | 0.62 | 0 | 1 | ERR349727 |
| PDB13-17_L1MF_Ch_H | Chad | Human | NUC | 23,046,238 | 15,617,912 | 67.77 | 15.47 | 13 | 3 | ERR349728,ERR563507,ERR563518 |
| PDB13-19_adF_Ch_D | Chad | Dog | NUC | 88,741,138 | 76,943,100 | 86.71 | 75.18 | 65 | 5 | ERR349731,ERR563509,ERR563520,ERR563533,ERR563544 |
| PDB13-27b1_adF_Ch_D | Chad | Dog | NUC | 139,764,890 | 95,458,521 | 68.3 | 92.70 | 74 | 5 | ERR349732,ERR563510,ERR563521,ERR563534,ERR563545 |
| PDB13-27c1_adF_Ch_D | Chad | Dog | MIT | 2,157,112 | 782,842 | 36.29 | 1.13 | 1 | 1 | ERR349733 |
| PDB13-27c1_L1MF_Ch_D | Chad | Dog | MIT | 15,484,644 | 10,348,781 | 66.83 | 10.26 | 8 | 3 | ERR349734,ERR563511,ERR563522 |
| PDB13-36_AdFem_Eth | Ethiopia | Human | NUC | 100,290,004 | 66,446,625 | 66.25 | 64.50 | 55 | 5 | ERR349735,ERR563512,ERR563523,ERR563535,ERR563546 |
| PDB13-36_L1mix_Eth | Ethiopia | Human | MIT | 1,023,952 | 637,308 | 62.24 | 0.92 | 1 | 1 | ERR349736 |
| PDB13-38_AdFem_Eth | Ethiopia | Human | MIT | 2,162,982 | 1,208,094 | 55.85 | 1.75 | 1 | 1 | ERR349737 |
| PDB13-38_L1mix_Eth | Ethiopia | Human | MIT | 10,231,108 | 5,883,801 | 57.51 | 5.87 | 5 | 3 | ERR349738,ERR563513,ERR563524 |
| PDB13-46 | Chad | Cat | NUC | 59,685,340 | 51,250,126 | 85.87 | 50.15 | 43 | 5 | ERR460386,ERR563548,ERR563554,ERR563561,ERR563566 |
| PDB13-78-Ad | Ethiopia | Baboon | NUC | 131,493,342 | 94,794,894 | 72.09 | 92.26 | 77 | 3 | ERR460387,ERR563549,ERR563555 |
| PDB13-78-L1 | Ethiopia | Baboon | NUC | 245,165,212 | 139,723,590 | 56.99 | 135.99 | 121 | 3 | ERR460389,ERR563550,ERR563556 |
| PDB13-92-Ad | Ethiopia | Dog | NUC | 59,912,414 | 52,943,803 | 88.37 | 51.67 | 43 | 5 | ERR460388,ERR563527,ERR563538,ERR563562,ERR563567 |
| PDB13-92-L1 | Ethiopia | Dog | NUC | 122,349,456 | 43,334,484 | 35.42 | 42.10 | 37 | 5 | ERR460390,ERR563551,ERR563557,ERR563563,ERR563568 |
| PDB14-100CH | Chad | Human | MIT | 8,469,766 | 6,871,657 | 81.13 | 8.32 | 7 | 2 | ERR1081332,ERR1243213 |
| PDB14-135CH | Chad | Human | NUC | 17,636,172 | 12,076,788 | 68.48 | 14.66 | 13 | 2 | ERR1081331,ERR1243212 |
| PDB14-207CH | Chad | Human | NONE | 219,034 | 1,666 | 0.76 | 0.00 | 0 | 2 | ERR1081362 |
| PDB14-209CH | Chad | Human | NONE | 465,120 | 15,711 | 3.38 | 0.02 | 0 | 1 | ERR1081322 |
| **PDB14-210CH** | Chad | Human | NONE | 822,112 | 1,090 | 0.13 | 0.00 | 0 | 3 | ERR1081361 |
| **PDB14-22CH** | Chad | Human | NONE | 377,076 | 1,623 | 0.43 | 0.00 | 0 | 1 | ERR1081336 |
| **PDB14-223CH** | Chad | Human | NUC | 12,766,930 | 11,176,393 | 87.54 | 13.55 | 11 | 2 | ERR1081325,ERR1243208 |
| **PDB14-24CH** | Chad | Human | NUC | 26,045,970 | 15,618,401 | 59.96 | 18.88 | 16 | 2 | ERR1081319,ERR1243206 |
| **PDB14-240MH** | Mali | Human | MIT | 374,388 | 261,532 | 69.86 | 0.38 | 0 | 1 | ERR1081341 |
| **PDB14-253MH** | Mali | Human | MIT | 608,986 | 296,201 | 48.64 | 0.43 | 0 | 1 | ERR1081340 |
| **PDB14-269MH** | Mali | Human | NONE | 599,264 | 376,797 | 62.88 | 0.54 | 0 | 1 | ERR1081338 |
| **PDB14-278EH** | Ethiopia | Human | MIT | 834,286 | 82,418 | 9.88 | 0.12 | 0 | 1 | ERR1081337 |
| **PDB14-279CH** | Chad | Human | NONE | 1,091,738 | 7,128 | 0.65 | 0.01 | 0 | 1 | ERR1081324 |
| **PDB14-283CH** | Chad | Human | MIT | 778,050 | 339,157 | 43.59 | 0.49 | 0 | 1 | ERR1081323 |
| **PDB14-68CH** | Chad | Human | MIT | 286,718 | 184,537 | 64.36 | 0.27 | 0 | 1 | ERR1081333 |
| **PDB14-69CH** | Chad | Human | NONE | 987,276 | 132,848 | 13.46 | 0.19 | 0 | 1 | ERR1081364 |
| **PDB15-18CH** | Chad | Human | MIT | 17,468,312 | 10,416,113 | 59.63 | 12.57 | 10 | 2 | ERR1081360,ERR1243224 |
| **PDB15-24CH** | Chad | Human | NONE | 306,614 | 97,299 | 31.73 | 0.14 | 0 | 1 | ERR1081330 |
| **PDB15-46CH** | Chad | Human | NONE | 151,762 | 2,543 | 1.68 | 0.00 | 0 | 1 | ERR1081329 |
| **REF** | Ghana | Human | NUC | 300,273,364 | 284,679,336 | 94.81 | 274.25 | 244 | 1 | ERR066175 |
| **ferret A female** | N/A* | N/A | N/A | 323,962,030 | 91,896,737 | 90.1 | 424.30 | 391 | 1 | ERR1945309 |
| **ferret B male** | N/A* | N/A | N/A | 316,685,164 | 274,866,059 | 86.79 | 399.50 | 365 | 1 | ERR1945310 |
| **ferret B female** | N/A* | N/A | N/A | 293,016,784 | 261,298,996 | 89.18 | 379.81 | 346 | 1 | ERR1945311 |

\*parasites originated from an infected dog from Chad

**Supplementary Table 2.** Details of *Dracunculus insignis* and *D. lutrae* samples, sequencing data and libraries used in this study. Reads and mapping statistics are for the sum across all sequenced libraries and lanes. ENA = European Nucleotide Archive.

| sample name | country | host | species | total reads | reads mapping | percent mapping | mean coverage | median coverage | number of lanes | ENA accession numbers |
| --- | --- | --- | --- | --- | --- | --- | --- | --- | --- | --- |
| Din88-31 | USA |  | insignis | 251825454 | 56729856 | 22.5 | 33.37 | 12 | 4 | ERR273919, ERR273941, ERR563496, ERR563502 |
| Din88-31L1 | USA |  | insignis | 147069250 | 32767530 | 22.3 | 21.43 | 8 | 4 | ERR273922, ERR273944, ERR563552, ERR563558 |
| DIut | Canada |  | lutrae | 87800942 | 2292804 | 2.6 | 2.14 | 1 | 2 | ERR1081356, ERR1243220 |

**Supplementary Table 3.** influence of different prior distribution assumptions on results of coalescence analysis. Values are means of the posterior distribution and 95% highest posterior density confidence intervals. Θ values are effective population size estimates and ! are divergence time estimates. East+Chad and Africa representing the two ancestral populations, as shown on Figure 5.

| prior | $\Theta_{W. AFRICA}$ | $\Theta_{CHAD}$ | $\Theta_{E. AFRICA}$ | $\Theta_{EAST+CHAD}$ | $\Theta_{AFRICA}$ | $\tau_{EAST+CHAD}$ | $\tau_{AFRICA}$ |
| --- | --- | --- | --- | --- | --- | --- | --- |
| $\alpha=10^{-8}$ | 21,538<br>(18,888-24,184) | 31,087<br>(20,496-40,550) | 7,849<br>(5024-10502) | 336,776<br>(251697-426576) | 112,057<br>(106969-117231) | 4,106<br>(2572-5542) | 20,643<br>(18326-22889) |
| $\alpha=10^{-7}$ | 21,625<br>(19160-24188) | 28,361<br>(18744-36941) | 7,151<br>(4756-9323) | 349,031<br>(252008-463045) | 112,494<br>(107749-117749) | 3,729<br>(2347-4809) | 20,760<br>(18660-22996) |
| $\alpha=10^{-6}$ | 21,147<br>(18633-23594) | 28,192<br>(18103-36990) | 7,370<br>(4603-9464) | 394,842<br>(273224-504162) | 112,903<br>(108006-118058) | 3,841<br>(2358-5080) | 20,398<br>(18047-22869) |
| $\alpha=10^{-5}$ | 21,563<br>(18831-24496) | 29,894<br>(20177-39670) | 7,617<br>(4998-10102) | 350,866<br>(253046-471265) | 112,844<br>(107483-118213) | 4,014<br>(2572-5421) | 20,612<br>(18150-23222) |
| $\alpha=10^{-4}$ | 21,233<br>(18420-23886) | 32,283<br>(22994-41750) | 8,153<br>(5596-10490) | 340,002<br>(244403-437895) | 112,930<br>(107743-118174) | 4,305<br>(2895-5559) | 20,319<br>(17666-22705) |

